# PoreMeth2: decoding the evolution of methylome alterations with Nanopore sequencing

**DOI:** 10.1101/2024.10.03.616449

**Authors:** Gianluca Mattei, Marta Baragli, Barbara Gega, Alessandra Mingrino, Martina Chieca, Tommaso Ducci, Gianmaria Frigè, Luca Mazzarella, Romina D’Aurizio, Francesco De Logu, Romina Nassini, Pier Giuseppe Pelicci, Alberto Magi

## Abstract

In epigenetic analysis, identifying differentially methylated regions (DMRs) typically involves detecting groups of consecutive CpGs that show significant changes in their average methylation levels. However, the methylation state of a genomic region can also be characterized by a mixture of patterns (epialleles) with variable frequencies, and the relative proportions of such patterns can provide insights into its mechanisms of formation.

Traditional methods based on bisulfite conversion and NGS, due to the read size (150 bp), allow epiallele frequency analysis only in high-CpG-density regions, limiting differential methylation studies to just 50% of the human methylome. Nanopore sequencing, with its long reads, enables the analysis of epiallele frequency across both high- and low-CpG-density regions.

We introduce a novel computational approach, PoreMeth2, an R library that integrates epiallelic diversity and methylation frequency changes from Nanopore data to identify DMRs, assess their formation mechanisms, and annotate them to genic and regulatory elements. We applied PoreMeth2 to cancer and glial cell datasets, demonstrating its ability to distinguish epigenomic changes with a strong effect on gene expression from those with a weaker impact on transcriptional activity.

PoreMeth2 is publicly available at https://github.com/Lab-CoMBINE/PoreMeth2.

## Background

In epigenetic analysis, a consolidated approach to detect methylation alterations between two samples consists in searching for groups of consecutive CpGs that concordantly show an increase (hyper-methylation) or a decrease (hypo-methylation) in their average methylation level (DMRs).

Currently used assays to determine CpGs methylation state are based on bisulfite conversion of methylated cytosines to uracil followed by SGS - RRBS and WGBS - which only allow characterisation of high CpG density regions (> 2 − 3 CpG/100 bp, representing 50% of the genome at best).

However, the last decade has seen the emergence of third-generation sequencing technologies, based on Nanopore sequencing [1], which allow to produce sequences in the order of tens to hundreds of kilobases (kb) and to directly recognize base modifications, such as 5mC, thus allowing concomitant analyses of genomic and epigenomic changes [2, 3, 4].

Using Nanopore and a novel computational method we recently reported that it is possible to infer the methylation state of 99% of all the CpG sites of the human genome (28.3 millions), with an average CpG density of 1 CpG/100bp, thus obtaining an unprecedented resolution for the identification of differentially methylated regions (DMRs) in low CpG density regions [5]. Most notably, application of this new technology to a chemoresistant leukemia dataset, allowed the identification of thousands of DMRs for each sample pair, with around 50% of them falling within low CpG density regions (≤ 2 CpG/100 bp), which are not detected by classical bisulphite-based methods [6].

Results of our analyses were highly informative for the mechanisms of drug-resistance in AMLs, but also confirmed previous studies [7] showing that a significant proportion of differentially-methylated genes were not differentially-expressed. Such results suggest that a large fraction of the DMRs observed in our samples may be merely passenger events that accompany cancer evolution with weak or no effect on gene expression [8].

The methylation state of a given genomic region (a group of adjacent CpG sites) in a cell population is defined not just by its average methylation level but by a mixture of patterns (epialleles) with variable frequencies. The relative proportion of such patterns can provide information on DMRs’ origin: an increase in the frequency of a specific epiallele suggests a selective formation, as opposed to multiple stochastic changes in the frequencies of many epialleles which are associated with random formation.

To date, WGBS, RRBS and methylation arrays have been used to study DMRs and epiallele composition. However, around 42% of 3-CpGs (58% of 4- and 70% of 5-CpGs) epialleles are larger than 150 bp, allowing the analysis of epiallele frequencies only in high-density CpG regions (CpG Islands, CGIs), strongly limiting their use in low-density CpG regions (< 3 CpG/100 bp) where short reads (150 bp) can overlap no more than two CpG sites. Long reads generated by Nanopore sequencing however reach lengths in the order of tens of Kb and are thus suitable to calculate epiallele frequency in both high- and low-density CpG regions, revolutionizing our capability to study methylome alterations.

In this work we introduce a novel computational approach that, combining epiallelic diversity changes with methylation frequency changes from Nanopore data, is capable of identifying DMRs and evaluate their mechanism of formation. The new approach was packaged in an R library, PoreMeth2, that also allows automatic annotation of DMRs with a new and efficient annotation scheme and generate useful graphical representations of the results.

We applied PoreMeth2 to a cancer dataset and a dataset of human peripheral glial cells (HPGC) treated with a G protein-coupled receptor agonist and we showed that our approach is capable to discriminate epigenomic alterations originated from selection of epialleles that have a stronger effect on gene expression from those generated by random rearrangement of epialleles with weaker effect on transcriptional activity.

## Results

### DMRs detection

To estimate the impact of read length on the analysis of epiallele diversity, we simulated reads of various sizes (100bp to 10 kb) and we evaluated their coverage across the human reference genome’s CpG dinucleotide map (hg19, see Methods). As depicted in Figure S1 of Supplemental Material, reads exceeding 5 kb enabled epiallele diversity assessment in at least 99% of the epigenome.

Long reads generated by Nanopore sequencing reach lengths in the order of tens of Kb, thus, are capable to study epiallele frequencies in both high- and low-density CpG regions revolutionizing our capability to study methylome alterations.

At present, the methylation state of a CpG site is studied by using the methylation frequency *β* (calculated as the ratio between the total number of CpG sites predicted as methylated and the total number of reads aligned to that CpG) and differential methylation between Test (T) and Control (C) samples by using Δ*β* = *β*_*T*_ −*β*_*C*_. Δ*β* takes values in the range [-1,1], where Δ*β* > 0 or < 0 indicates, respectively, hyper- or hypo-methylation of the Test vs. Control samples.

Recently, we developed a novel tool (PoreMeth) based on a heterogeneous form of the shifting level model (SLM) that is capable of identifying DMRs by segmenting methylation frequency differences (Δ*β*) inferred from Nanopore data [5].

In this work we expand the PoreMeth tool to include epiallelic diversity changes. The diversity of DNA methylation patterns in a cell population can be measured by using the Shannon entropy [9]:

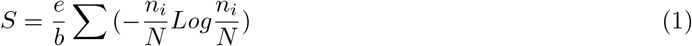

where *e* is entropy for code bit, *b* is the number of CpG sites, *n*_*i*_ is the occurrence of methylation pattern *i* and *N* is the total number of reads overlapping the *b* CpG sites. For each genomic feature, S was calculated by using *b* = 3 and averaged across all the CpG sites within the feature.

DNA-methylation entropy takes values in the range [0, 1] and is 0 when all cells share same DNA-methylation patterns, and 1 when instead all possible patterns are equally represented (Figure 1.a). Studying the differential entropy of a DMR (Δ*S*) can thus allow to distinguish between the selection of a specific epiallele (Δ*S* < 0) from the stochastic changes of multiple epialleles (Δ*S* > 0).

**Figure 1:**
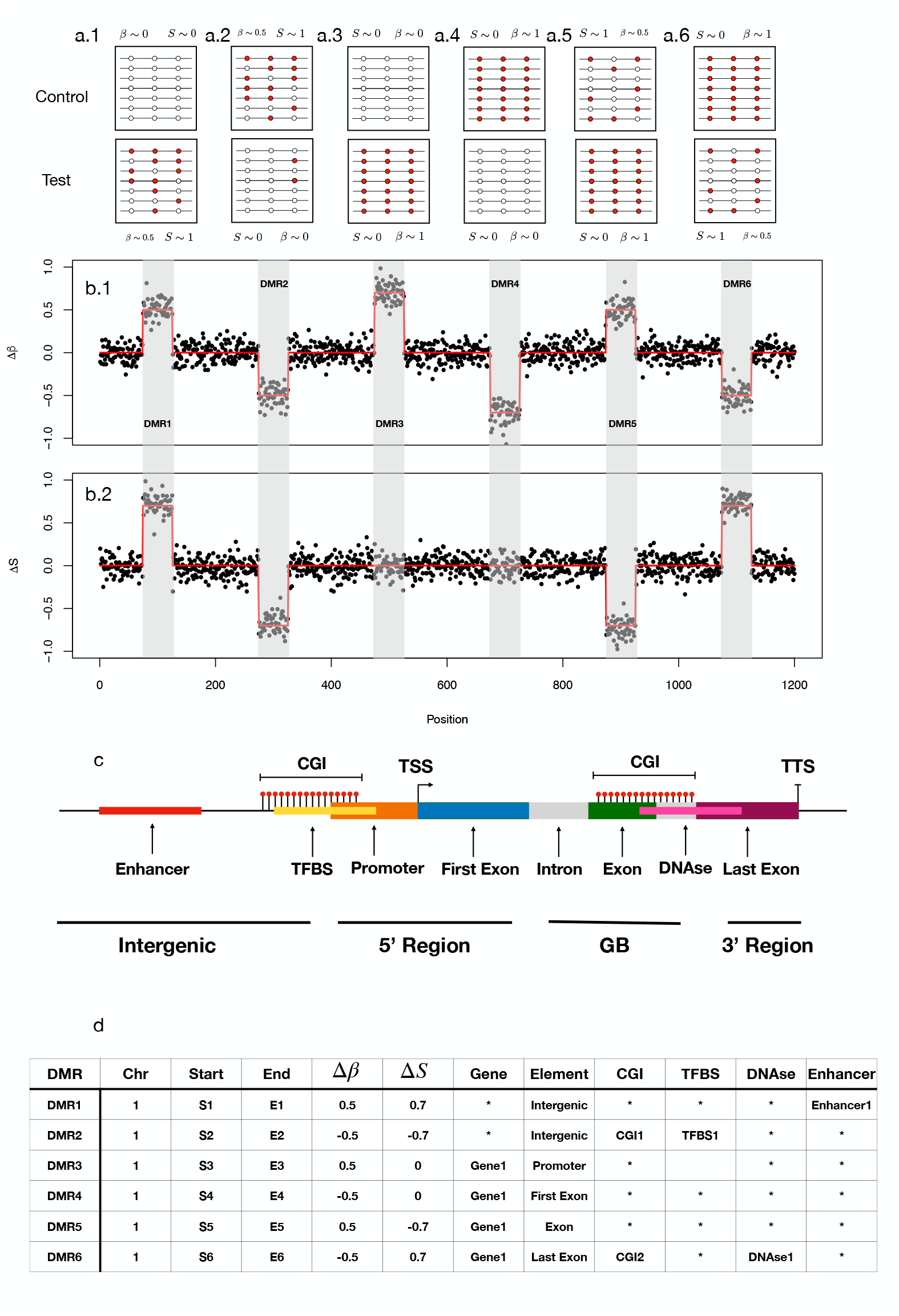
Computational workflow of PoreMeth2. Panel (a) shows a schematic representation of the six possible epiallelic changes between test and control samples: hyper-methylation with entropy increase (Δ*S* > 0.1 and Δ*β* > 0,a1), hypo-methylation with entropy decrease (Δ*S* < 0 and Δ*β* > 0, a2), hyper- and hypo-methylation with no entropy change (Δ*S* ∼ 0 and Δ*β* > 0,a3, Δ*S* ∼ 0 and Δ*β* < 0,a4), hyper-methylation with entropy decrease (Δ*S* < −0.1 and Δ*β* > 0,a5) and hypo-methylation with entropy increase (Δ*S* > 0 and Δ*β* < 0, a6). PoreMeth2 takes as input the methylation calls from Nanopolish or Guppy and calculates methylation frequency and entropy. Panel (b) shows Δ*β* (b.1) and Δ*S* (b.2) signals calculated for each CpG dinucleotide and ordered for genomic position. The two signals show six DMRs that reflect the epiallelic diversity changes reported in panel (a). In order to identify epiallelic composition changes the two signals *x*_*i*_ are modeled with SLM as the sum of two independent stochastic processes (*x*_*i*_ = *m*_*i*_ + *ϵ*_*i*_), where *m*_*i*_ = (*m*_*i*_1, *m*_*i*_2) is the vector of the unobserved mean level and *ϵ*_*i*_ is the vector of white noises. The white noise vector *ϵ*_*i*_ follows a bivariate normal distribution with mean *µ*_*ϵ*_ = [0] and covariance matrix Σ_*ϵ*_; *z*_*i*_ are random variables taking the values in [0,1] with probabilities *η* = *Pr*(*z*_*i*_ = 1) (1 − *η* = *Pr*(*z*_*i*_ = 0)); *δ*_*i*_ are random vectors that follow a bivariate normal distribution and*µ*_*i*_ is the vector of the means (see methods). DMRs identified with the bivariate version of the SLM algorithm can then be annotated with a scheme that reports all the genic elements overlapping a DMR, and for each of these element it calculates the overlap with regulatory feature (CGI, Enhancers, TFBS, and DHS). Panel (c) shows the gene model used for PoreMeth2 annotation and panel (d) the annotation results of the six DMRs.

In particular, by combining Δ*S* and Δ*β* we can observe six different possible epiallelic diversity changes between test and control samples (Figure 1.a): stochastic change with hyper-methylation (Δ*S* > 0 and Δ*β* > 0, Figure 1.a.1) and hypo-methylation (Δ*S* > 0 and Δ*β* < 0, Figure 1.a.6), selective change with hyper-methylation (Δ*S* < 0 or Δ*S* ∼ 0 and Δ*β* > 0, Figure 1.a.3 and Figure 1.a.5) and selective change with hypo-methylation (< 0 or Δ*S* ∼ 0 and Δ*β* < 0, Figure 1.a.2 and a.4).

In order to identify selective or stochastic epiallelic changes, we developed a bi-variate version of the SLM algorithm (BiSLM) that is capable to simultaneously analyze and segment Δ*S* and Δ*β* values (see Methods). In summary, our method processes the Δ*S* and Δ*β* values of consecutive CpG dinucleotides to identify genomic regions exhibiting increased or decreased methylation and entropy levels between two samples (Figure 1.b).

In order to evaluate the ability of our algorithm to identify DMRs of different size and with different epiallelic changes we applied BiSLM on simulated synthetic methylation profiles and we found that it is capable to detect DMRs as small as five consecutive CpGs with sequencing coverage greater than or equal to 20x (see Supplemental Material and Figure S8 of Supplemental Material).

### DMRs annotation

A number of studies have shown that methylation of promoters in CGIs leads to the expression down-regulation of tumor suppressor genes, thus representing a critical mechanism in cancer development [10]. Recently we demonstrated that hyper-methylated genes at sparse CpGs in the gene body are significantly enriched in transcription factors (TFs) that deregulate large gene regulatory networks inducing drug resistance in AML patients [5]. Furthermore, other studies demonstrated that DNA methylation at transcription factor binding sites (TFBS) can influence gene expression by regulating the ability of transcription factors to bind to their target DNA sequences [11].

These results demonstrate the fundamental importance of studying the overlap between genic elements (promoters, introns, exons) and regulatory features (CGIs, TFBS, enhancers) affected by DMRs in order to elucidate their impact on gene expression and phenotypes.

At present, few tools have been developed for annotating genomic intervals to genic and regulatory elements, among which GoldMine [12], annotatePeaks.pl from Homer tool [13], GenomicDistributions [14] and the R package annotatr [15]. These tools can annotate genomic intervals by following one of two different strategies: i) reporting a single genic feature by using feature priority (using gene models with the priority order promoter > 3 end > exon > intron > intergenic) or ii) reporting each genic and regulatory feature overlapping the interval as a row (long format).

The feature priority annotation scheme allows to obtain only partial information on the functional impact that a DMR may generate, especially when the epigenomic alterations are large and affect multiple genes and multiple regulatory features, while the ‘all feature’ scheme is very complex to be summarized. Moreover, none of these methods allow to study the reciprocal overlap between gene model elements and regulatory feature, thus limiting the interpretation of functional effect of a DMR.

For these reasons, we implemented a novel annotation scheme that reports all the genic elements (Promoter, First Exon, internal introns and exons and 3’UTR) overlapping a DMR, and for each of these elements, it calculates, when present, the overlap with regulatory features such as CGI, Enhancers, TFBS, and DNase I hypersensitive sites (DHS) (see Figure 1.c).

This annotation scheme not only identifies each genic element affected by a DMRs, but it also allows to evaluate its functional interaction with regulatory features. Moreover, each overlap is quantified in terms of percentage allowing to discriminate genomic elements where few bases are affected by a DMR from those where the DMRs have a greater overlap, permitting a more precise interpretation of its functional impact.

The BiSLM algorithm and the novel annotation scheme were integrated in a R package named PoreMeth2 that allows to automatically identify and annotate DMRs by comparing the Nanopore methylation data of a pair of test and matched normal samples (see Methods). PoreMeth2 also allows a gene-based annotation using feature priority scheme, in which each gene affected by a DMR is classified as either 5’ regulatory regions (5’Reg, if the DMR overlaps with the promoter, 5’UTR, or the first exon), 3’ untranslated regions (3’UTR, if the DMR overlaps with the last exon but not with 5’Reg elements) or gene bodies (GB, if the DMR overlaps with internal introns or exons but not with 5’Reg or 3’UTR elements). The feature priority scheme maintains the regulatory elements overlap, and can be very useful to study the correlation of DMRs with other omic layers such as gene expression. The Annotation functions of PoreMeth2 are powered by fortran libraries that allow to annotate tens of thousands of DMRs in parallel in minutes (Figure 1.c and Methods).

### AML data analyses

To test the power of PoreMeth2 we analysed methylation data from AML sample pairs that we previously analyzed in [5], in which we demonstrated that in relapsed AML hyper-methylated genes at sparse CpGs in the gene body, were significantly enriched in cancer genes (Oncogenes and Tumor Suppressor) and cancer-related pathways. The dataset consists of sample pairs at diagnosis (T) and relapse (R) from three AML patients (UD5, UD10 and AML2) who received standard chemotherapy and relapsed with chemoresistant disease (see methods). The six samples were sequenced with ONT sequencer obtaining, for each sample, a sequencing coverage of 20-30x (see Methods and Supplemental Material).

As a first step we calculated methylation frequency (*β*) and entropy (*S*) for each sample, we applied BiSLM to each pair of AML samples (see Methods) and we classified DMRs in six different categories reflecting Δ*β* (hyper-Methylation, Δ*β* > 0.2 and hypo-Methylation, Δ*β* < 0. −2) and Δ*S* (hyper-entropic, Δ*S* > 0.1, iso-entropic, −0.1 < Δ*S* < 0.1, and hypo-entropic, Δ*S* < − 0.1) variations. Due to the read size obtained by our Nanopore runs, we were able to calculate epiallelic diversity measures (*S*) for 80-90% of epialleles with at least 5 reads (Figure S9 of Supplemental Material).

Our algorithm identified 3102 DMRs for UD5 (3.18 Mb of genomic regions), 2825 for UD10 (3.07 Mb) and 2874 for AML2 (3.18 Mb), with a significantly larger fraction of hyper-methylated vs. hypo-methylated DMRs, both in terms of numbers (1999 vs 1103 for UD5, 2292 vs 533 for UD10 and 2575 vs 299 for AML2, Figure 2.a) and total size (1930 kb vs 1250 kb for UD5, 2570 kb vs 500 kb for UD10 and 2860 kb vs 320 kb for AML2, Figure S10 Supplemental Material). Moreover, both hyper and hypo-methylated DMRs were equally distributed among the three epiallelic change categories in terms of number (Figure 2.a) and size (Figure S10 of Supplemental Material).

**Figure 2:**
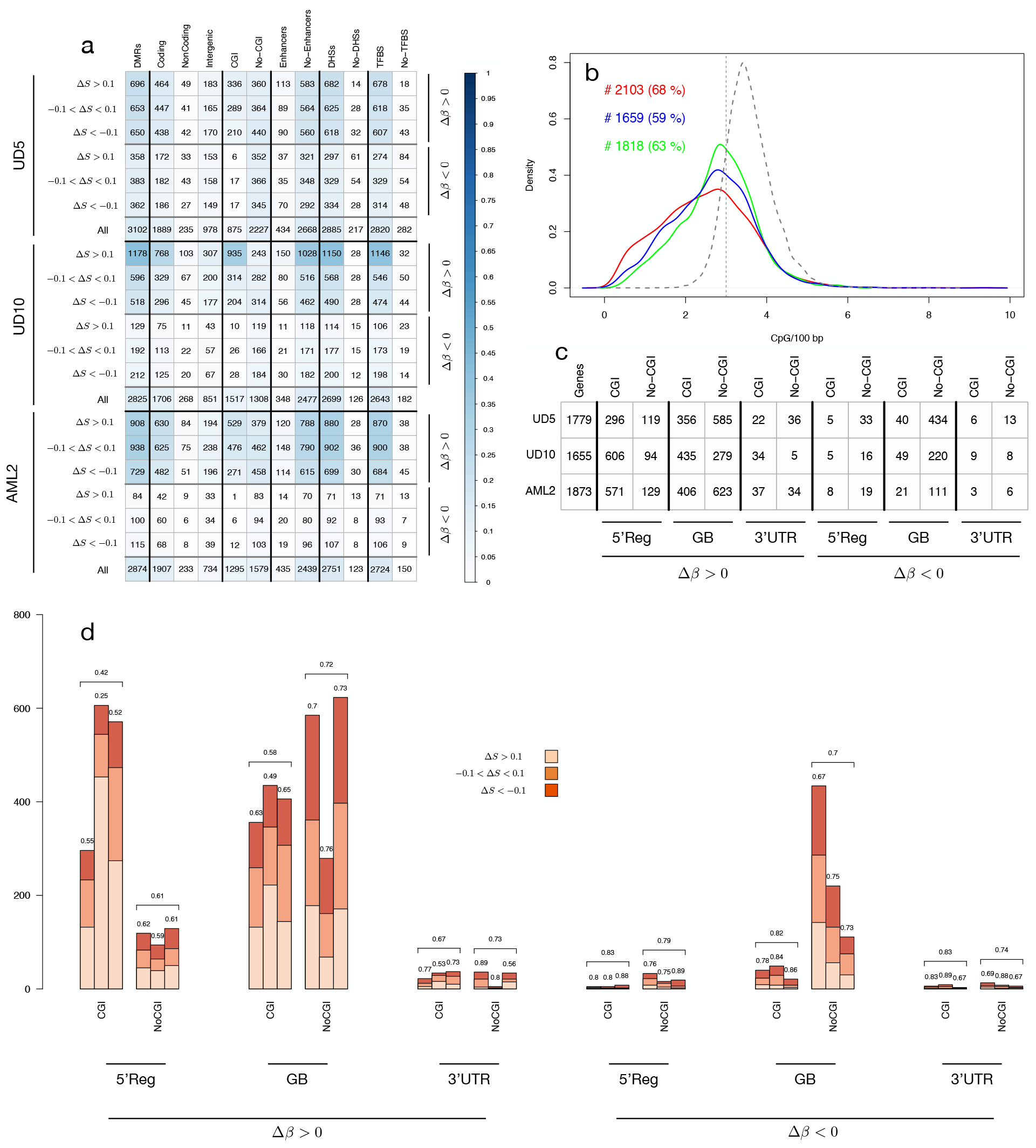
DMRs of the three pairs of AML samples. Panel (a) shows the number of DMRs detected by BiSLM in the three AML pairs for all the six categories reflecting Δ*β* (hyper-Methylation, Δ*β* > 0.2 and hypo-Methylation, Δ*β* < −0.2) and Δ*S* (hyper-entropic, Δ*S* > 0.1, iso-entropic, −0.1 < Δ*S*¡0.1, and hypo-entropic, Δ*S* < −0.1) variations. Numbers are reported for DMRs overlapping protein coding genes (Coding), non coding genes (non-coding), intergenic regions (Intergenic), CpG islands (CGI), Enhancers, DNase I hypersensitive sites (DHSs) and transcription factor binding sites (TFBS). NoCGI, No-Enhancers, No-DHSs and No-TFBS represent the number of DMRs that do not overlap with CGI, Enhancers, DHSs and TFBS respectively. The color intensity in each cell reflects the proportion of DMRs of a category with respect to all DMRs for each sample according to colorbar. Panel (b) shows CpG density distribution of the DMRs detected by BiSLM on the three pairs of AML samples. Vertical continuous line indicates the resolution limit of WGBS (≤ 2 CpG/100 bp), while vertical dotted line indicates ERRBS resolution limit (≤ 3 CpG/100 bp). Text on left (right) side of the the plot reports the total number (#) and percentage (%) of DMRs detected by PoreMeth with CpG density ≤ 2 CpG/100 bp (≤ 3 CpG/100 bp). The table of (c) reports the total number of DM genes with DMRs at different genomic elements (5’ regulatory region, Reg, internal introns and exons, GB, 3’UTR) inside CpG islands (CGI) and outside CpG islands (NoCGI). The barplot of panel (d) reports the number of hyper-entropic (Δ*S* > 0.1), hypo-entropic (Δ*S* < −0.1) and iso-entropic (−0.1 < Δ*S* < 0.1) epiallelic changes for hyper- and hypo-methylated genes with DMRs overlapping different genic elements at CGI and sparse CpGs (NoCGI). Numbers above bars show the percentage of hypo-entropic (Δ*S* < −0.1) or iso-entropic (−0.1 < Δ*S* < 0.1) DMGs. Horizontal brackets above each group of three bars summarize average percentages of the three samples.

Average size of DMRs was 700–800bp for all three AML pairs, with a distribution ranging from hundreds bp to tens kb and no significant differences between the six DMR categories (Figure S11 of Supplemental Material). Remarkably, ∼ 70% of DMRs identified by our method showed a CpG density ≤ 3 CpG/100bp (e.g. the resolution limit of standard NGS reads) demonstrating that long-read sequencing coupled to our novel computational method allows the identification of epiallelic changes between test and control samples at unprecedented resolution, extending analyses of high density CpG regions (CpG islands or CGIs), as achieved until now, to sparse CpGs (Figure 2.b).

To evaluate the functional impact of the epiallelic changes identified by our segmentation strategy, we used the annotation module of PoreMeth2 and we studied the distribution of DMRs across genic and regulatory elements. Most DMRs mapped within annotated protein coding genes (62, 67 and 66% for UD5, UD10 and AML2, respectively), around 10% in non-coding genes (Non-coding RNAs, pseudogenes and processed transcripts) and 30% in intergenic regions (non overlapping GenCode elements). Moreover, the great majority of DMRs overlapped DHSs (90, 92 and 93%) and TFBS (90, 92 and 93%), with a small fraction also overlapping enhancers (13, 12 and 13%) (Figure 2.a). As expected from CpG density distribution, 18, 44 and 29% DMRs (for UD5, UD10 and AML2 respectively) overlapped CGIs, while the remaining were located in low density CpG regions (Figure 2.a).

As a further step, we used the annotation module of PoreMeth2 to classify protein coding genes in three main functional categories: 5’ regulatory regions (5’Reg, if the DMR overlaps with promoter, 5’UTR or first exon), 3’untranslated regions (3’UTR, if the DMR overlaps with the last exon but not with Reg elements) and gene-bodies (GB, if the DMR overlaps with internal introns or exons but not with Reg or 3’UTR elements). The number of protein coding genes affected by DMRs (differentially-methylated genes, DMGs) were ∼ 1, 700 per patient (1779, 1655 and 1873 for UD5, UD10 and AML2 respectively, Figure 2.c), most of which hyper-methylated (∼ 70, ∼ 85 and ∼ 93% in UD5, UD10 and AML2, respectively, Figure 2.c). 25-40% showed DMRs at 5’Reg, 60-80% at GB and ∼ 5% at 3’UTR (Figure 2.c).

DMGs with hyper-methylated DMRs in 5’Reg mainly involved CGIs (∼ 82% across the three samples: 71, 86 and 81% for UD5, UD10 and AML2), while DMG with hyper-methylated DMRs at gene-bodies mostly overlapped sparse CpGs (∼ 55% across the three samples). Hypo-methylated genes were almost entirely associated with DMRs overlapping sparse CpGs (∼ 90%), regardless of their position within genes (Figure 2.d). Moreover, the great majority of DMGs showed hypo-entropic (Δ*S* < − 0.1) or iso-entropic (− 0.1 < Δ*S* < 0.1) epiallelic changes (between 60 and 80% for both hyper- and hypo-methylation), with the exception of DMGs with hyper-methylated DMRs in 5’ regulatory regions at CGIs (for which hyper-entropic changes represents 60%, Figure 2.d).

To investigate the impact of DMRs on gene expression, we analysed the six AML samples with triplicate RNA-sequencing experiments (RNAseq) and studied differential gene-expression between Relapse and Diagnosis samples using DESeq2 [16] (see Methods). We identified 3,997, 4,677 and 1,759 differentially-expressed genes (DEGs; Supplemental Data 2) in UD10, UD5 and AML2, respectively, with different ratios of over- and under-expressed genes (2,044 and 1,953 in UD5; 2,890 and 1,787 in UD10, 495 and 1,264 in AML2) (Table 4 of Supplemental Material).

As shown in Figure S12 of Supplemental Material, among all the DMGs, the proportion of DEGs is similar for genes hosting hyper-entropic (Δ*S* > 0.1) and hypo/iso-entropic (Δ*S* < 0.1) DMRs (10% for hyper-methylated genes on CGIs and 30% for other categories). Surprisingly, considering only DM-DEGs, we found that the great majority is affected by hypo/iso-entropic (Δ*S* < 0.1) epiallelic changes, suggesting that epiallelic selection have stronger effect on gene transcription (Figure 3.a). We then analysed the proportion of DMGs that were also DEGs (DM-DEGs) with respect to all DEGs considering separately genes affected by DMRs at different genic elements, in CGIs or sparse CpGs, and we found that mainly DMGs with DMRs at sparse CpGs in gene-body with Δ*S* < 0.1 (hypo- and iso-entropic) are significantly enriched of DEGs across the three samples (Figure 3.b).

**Figure 3:**
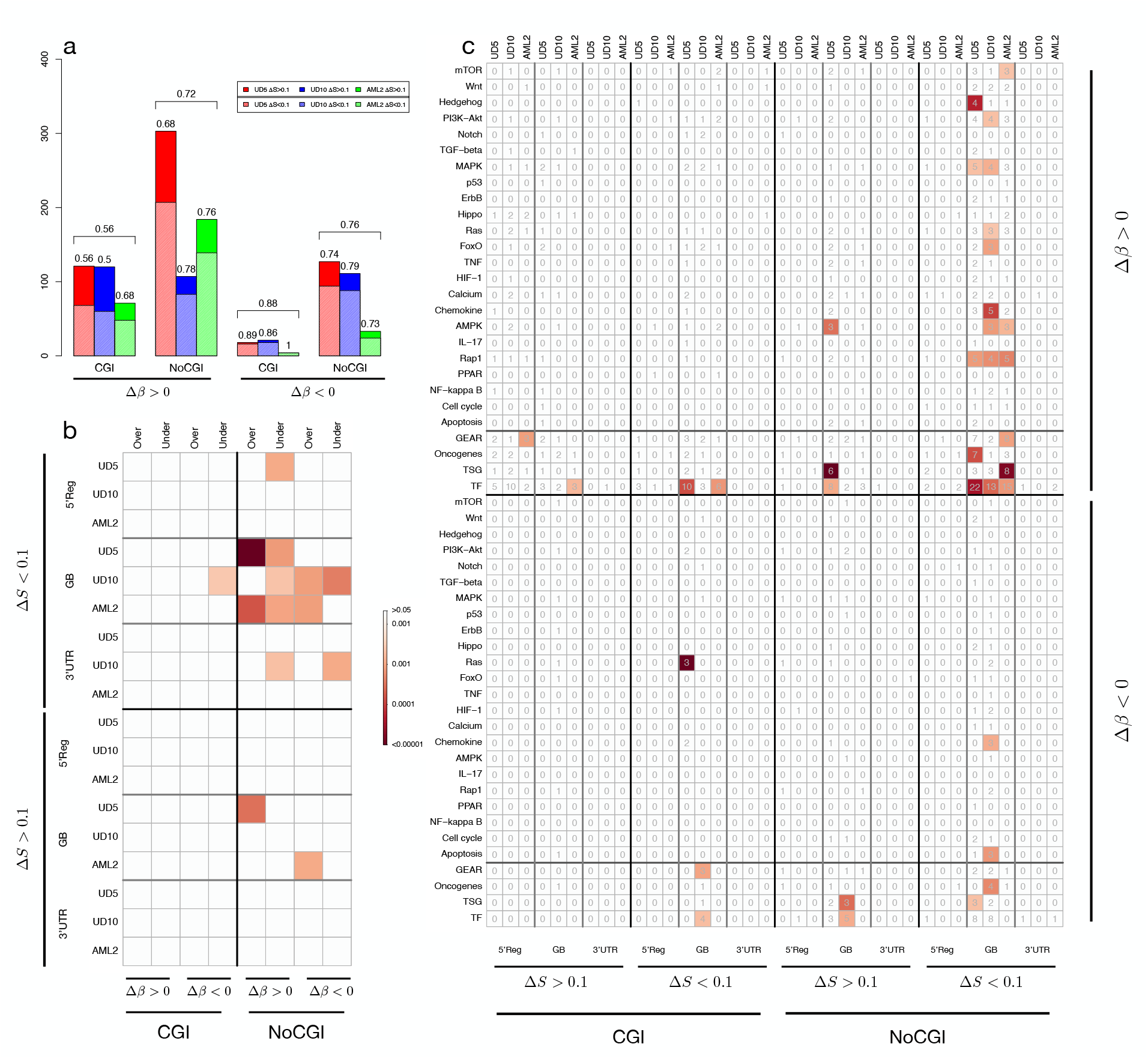
DMRs and differential expression. Panel (a) shows the proportion of DM-DEGs with hyper-entropic (Δ*S* > 0.1) and iso- or hypo-entropic (Δ*S* < 0.1) DMRs. Results are reported for hyper- (Δ*β* > 0.2) and hypo-methylated (Δ*β* < −0.2) DMRs overlapping CpG islands (CGI) and outside CpG islands (NoCGI). Textured bars show the number of DM-DEGs with iso- or hypo-entropic (Δ*S* < 0.1) DMRs. Panel (b) shows the results of ORA for DM-DEGs. The analysis was performed separately for genes affected by DMRs at the 5’ regulatory region (5’Reg), the gene body (GB) and the 3’UTR and for DMRs overlapping CpG islands (CGI) and outside CpG islands (NoCGI). The blocks of the corrplot reports the results of ORA for different classes of DMRs formation: Δ*β* > 0.2 and Δ*S* < 0.1 (hyper-methylated and hypo- and Iso-entropic), Δ*β* > 0.2 and Δ*S* > 0.1 (hyper-methylated and hypo- and Iso-entropic), Δ*β* < −0.2 and Δ*S* < 0.1 (hypo-methylated and hypo- and Iso-entropic), Δ*β* < −0.2 and Δ*S* > 0.1 (hypo-methylated and hypo- and Iso-entropic). The color intensity in each cell reflects statistical significance according to colorbar. For each category, Fisher exact tests were calculated comparing the number of DM-DEGs, DMGs, DEGs and all genes tested in RNA-seq experiments. Panel (c) shows the results of ORA for under-expressed DM-DEGs. The analysis was performed separately for genes affected by DMRs at the 5’ regulatory region (5’Reg), the gene body (GB) and the 3’UTR and for DMRs overlapping CpG islands (CGI) and outside CpG islands (NoCGI). The blocks of the corrplot reports the results of ORA for different classes of DMRs formation: Δ*β* > 0.2 and Δ*S* < 0.1 (hyper-methylated and hypo- and Iso-entropic), Δ*β* > 0.2 and Δ*S* > 0.1 (hyper-methylated and hypo- and Iso-entropic), Δ*β* < −0.2 and Δ*S* < 0.1 (hypo-methylated and hypo- and Iso-entropic), Δ*β* < −0.2 and Δ*S* > 0.1 (hypo-methylated and hypo- and iso-entropic). The numbers in each cell represent the total number of DM-DEGs for each category, while the color intensity reflects statistical significance according to colorbar. Fisher exact test and number of genes were calculated for cancer-related pathways selected from KEGG database, TFs, TSG and Oncogenes selected by COSMIC and GEAR genes.

As a final step, in order to evaluate DMRs’ effect in terms of expression at each genomic feature, we performed over-representation analyses (ORA) of DMGs pertaining to each subclass of DMRs, that were also DE. ORA was performed on DM-DE genes against a collection of cancer related pathways (KEGG, [17]), tumor suppressor genes (TSGs) and oncogenes (from COSMIC databes, [18]), transcription factors (TFs, [19]) and drugs resistance-associated genes (GEAR, [20]). DM-DE genes with DMRs overlapping CpG islands showed few significant over-representation of the tested datasets, regardless of methylation status or expression (Figure 3.c). The same was observed for the DM-DEGs at sparse CpGs with hyper-entropic epiallelic changes (Δ*S* > 0.1). hyper-methylated DM-genes with Δ*S* < 0.1 (hypo- or iso-entropic) at sparse CpGs in gene bodies, instead, were enriched in TFs and cancer pathways in all three patients (Figure 3.c).

Taken as a whole, these results demonstrate that hypo- or iso-entropic epiallelic changes (Δ*S* < 0.1) have a stronger impact on gene expression than changes generated randomly, and that these genes are enriched in cancer pathways.

### Human Peripheral Glial Cells

To test our method on a different experimental setup, suited to the evaluation of methylation state evolution, we applied the analysis to sequencing data obtained from HPGC cultures before (T0) and after 48 hours (T48) treatment with a G protein-coupled receptor (GPCR) agonist (See Methods and Supplemental Material).

Methylation frequency (*β*) and entropy (*S*) were calculated for T0 and T48 samples and we then applied BiSLM to classify DMRs in six different categories reflecting Δ*β* and Δ*S* variations (as in previous section). Given the high coverage and read size, we were able to calculate Δ*S* values for more than 99% of epialleles with at least 5 reads (Figure S13 of Supplemental Material).

BiSLM identified 636 DMRs, most of which are hyper-methylated in terms of both number and length (616 hyper-methylated vs 20 hypo-methylated, Figure 4.a). Additionally, 80-95% of the DMRs (80% for the hyper- and 95% for the hypo-methylated) are iso- or hypo-entropic (Figure 4.a). Cumulative size and size distribution for the six DMR categories are shown in Supplementary Figure 14 and Supplementary Figure 15 respectively.

**Figure 4:**
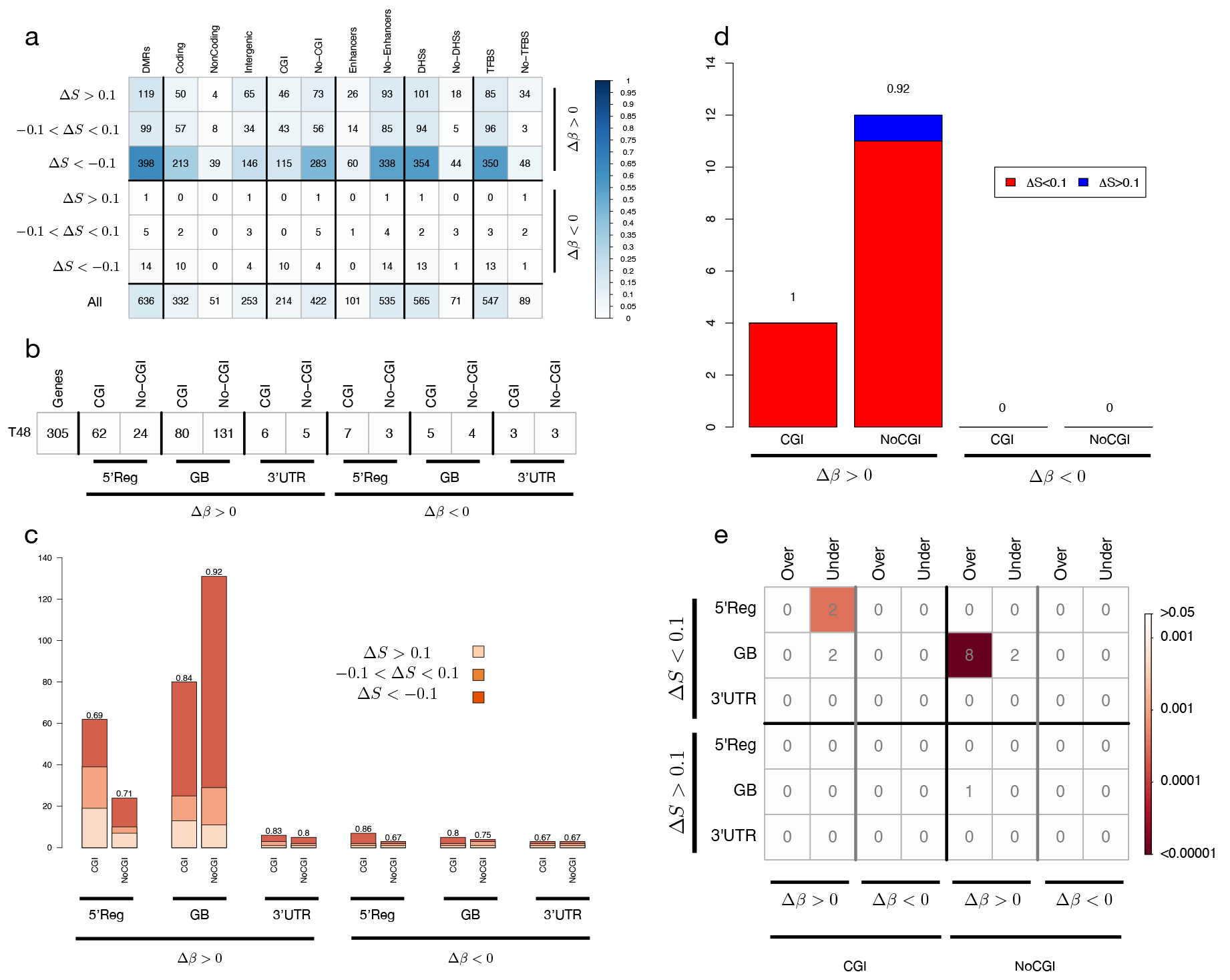
DMRs of the HPGC samples. Panel (a) shows the number of DMRs identified for each of the six categories defined by Δ*β* and Δ*S* values. The plot reports the number of DMRs overlapping each genic (coding, non coding and intergenic) and regulatory (CGI, Enhancer, DHS, TFBS) element class., while the color intensity in each cell reflects the proportion of DMRs of a category with respect to all DMRs according to colorbar. Panel (b) reports the number of DMGs affected by hyper-methylated (Δ*β* > 0.2) and hypo-methylated (Δ*β* < −0.2) DMRs overlapping different genic features (5’Reg, Gene Body and 3’UTR), outside and inside CpG Islands. The barplot in (c) displays the number of DMGs reported in (b), further categorized in hyper-entropic (Δ*S* > 0.1), iso-entropic (−0.1 < Δ*S* < 0.1) and hypo-entropic (Δ*S* < −0.1). Panel (d) shows the proportion of DM-DEGs with hyper-entropic (Δ*S* > 0.1) and iso- or hypo-entropic (Δ*S* < 0.1) DMRs. Results are reported for hyper- (Δ*β* > 0.2) and hypo-methylated (Δ*β* < −0.2) DMRs overlapping CpG islands (CGI) and outside CpG islands (NoCGI).

DMRs were then annotated with the annotation module of PoreMeth2 and we found that ∼ 50% mapped within annotated protein coding genes (332), ∼ 8% in non-coding genes and ∼ 40% in intergenic regions. As in previous section, we found that the great majority of DMRs overlapped DHSs (88%) and TFBS (86%), while only a small fraction with Enhancers (16%). As expected from DMRs’ CpG density distribution, only ∼ 34% mapped with CGIs, while the remaining ∼ 66% overlapped low-density CpG regions (Figure 4.a).

We then used the annotation module of PoreMeth2 to classify protein coding genes affected by DMRs (DMGs) in three main functional categories: 5’ regulatory regions (5’Reg), 3’untranslated regions (3’UTR) and gene-bodies (GB). The total number of DMGs is 305, nearly all of which have a hyper-methylated DMR in the 5’Reg or GB (86 in 5’Reg and 211 in GB, Figure 4.b). DMGs with hyper-methylated DMRs in 5’Reg mainly involved CGIs (62 in CGI vs 24 in NoCGI), while DMG with hyper-methylated DMRs at gene-bodies mostly overlapped sparse CpGs (131 in NoCGI vs 80 in CGIs). The vast majority of DMGs exhibit hypo-entropic (Δ*S* < − 0.1) or iso-entropic (− 0.1 < Δ*S* < 0.1) changes, especially those with DMRs in the GB within low-density CpG regions, where hypo-entropic DMRs account for nearly 80% (Figure 4.c).

To study the impact of methylation on gene expression, we conducted quadruplicate RNA-seq experiments for the T0 and T48 samples using Nanopore sequencing and then analyzed differential expression with DESeq2 (see Methods) and correlated with differential methylation. Only a small fraction of DMGs are also DEGs, and these consist solely of DMGs with hyper-methylated DMRs (Figure S16 of Supplemental Material). Remarkably, almost all DM-DEGs showed hypo-entropic (Δ*S* < − 0.1) or iso-entropic (− 0.1 < Δ*S* < 0.1) epiallelic changes (Figure 4.d).

Finally, we analyzed the proportion of DM-DEGs with respect to all DEGs, and we found that only DMGs with hypo- and iso-entropic DMRs (Δ*S* < 0.1, at 5’Reg in CGI and at GB in low density CpG regions) are significantly enriched in DEGs (Figure 4.e). These results further demonstrate that our new computational approach, through the use of differential entropy, can distinguish between DMRs that have a direct impact on gene expression and those that have a weak effect on transcriptional activity.

## Discussion and Conclusion

In this work we present the first computational method for the identification of DMRs and simultaneous prediction of their mechanism of origin from read-level methylation calls obtained with Nanopore sequencing of two samples - test and control. To this end we combined methylation frequency (Δ*β*) - to detect increases or decreases of methylation levels - with methylation entropy (Δ*S*) - to measure variations in epiallelic composition.

Our computational strategy consists in jointly segmenting Δ*β* and Δ*S* signals by means of a bivariate version of the SLM (BiSLM) algorithm to identify consecutive CpG dinucleotides that show increases or decreases in their mean values. Synthetic analyses demonstrated that our approach requires sequencing coverages larger than 20x to correctly identify DMRs with as few as five consecutive CpG and to predict their epiallelic change.

The BiSLM algorithm has been packaged in an R library (PoreMeth2) that also includes functions for DMRs annotation with respect to both genic and regulatory elements. The annotation function is capable of annotating all the genic elements overlapping a DMR, and to calculate the overlap with regulatory features for each genic element (such as CGI, Enhancers, TFBS and DHSs) thus allowing a better interpretation of the functional effect of methylation alterations.

To demonstrate the power of the PoreMeth2 pipeline, we first applied it to the analysis of three AML sample pairs at diagnosis (T) and relapse (R) that we previously analyzed in [5]. BiSLM identified around 3,000 DMRs for each pairs of samples with a significantly larger fraction of hyper-methylated vs. hypomethylated. In accordance with the results obtained in [5] approximately ∼ 70% of DMRs showed a CpG density of ≤ 2 CpG/100bp, demonstrating that long-read sequencing coupled to our novel computational method allows the identification of epiallelic changes at unprecedented resolution, extending analyses of high density CpG regions (CpG islands or CGIs), as achieved until now, to sparse CpGs.

As in [5], annotation of DMRs showed that the involvement of sparse CpGs was predominant in genes hyper-methylated at gene-bodies and that only DMGs with hyper-methylated DMRs at sparse CpGs in gene-body have a statistically significant impact on gene expression across the three samples. Remarkably, DMRs with Δ*S* < 0.1 (hypo- and iso-entropic) have the highest impact on gene expression, while hyper-entropic DMRs have marginal effect.

As a further step we used PoreMeth2 to analyze a Nanopore dataset of HPGC cells before and after treatment with GPCR agonist. BiSLM between treated and non-treated cells identified 636 DMRs, mostly hyper-methylated and with the majority being iso- or hypo-entropic.

About 50% of DMRs mapped to protein-coding genes, and most DMGs had hyper-methylated DMRs in 5’ regulatory regions or gene bodies. Only a small fraction of DMGs were also DEGs, and almost all DM-DEGs showed hypo-entropic (Δ*S* < − 0.1) or iso-entropic (0.1 < Δ*S* < 0.1) epiallelic changes, demonstrating that selected epialleles exert a significant effect on gene expression.

These results demonstrate that our approach allows us to discriminate epigenomic alterations originated from selection of epialleles that have a direct effect on gene expression from those generated by the random rearrangement of epialleles with low impact on gene expression. In conclusion, PoreMeth2 is the first computational pipeline that is capable of exploiting the intrinsic characteristics of long read methylation data to study methylation at an unprecedented resolution. Moreover, the data generated by ONT devices can also be applied to other DNA modifications, such as 5hmC and 6mA. At present, we are testing PoreMeth2 in the analysis of 5hmC profiles in liquid and solid cancers.

## Methods

### Bivariate SLM algorithm

SLM are a special class of Hidden Markov Models in which sequential observations *x* = (*x*_1_, …, *x*_*i*_, …, *x*_*N*_) are considered to be realizations of the sum of two independent stochastic processes *x*_*i*_ = *m*_*i*_ + *ϵ*_*i*_, where *m*_*i*_ is the unobserved mean level and *ϵ*_*i*_ is normally distributed white noise.

In order to jointly segment Δ*S* and Δ*β* values of consecutive GpG dinucleotide, we extended the classical SLM model to a bi-variate version where *x*_*i*_ = (Δ*S*_*i*_, Δ*β*_*i*_), *m*_*i*_ = (*m*_*i*1_, *m*_*i*2_) and *ϵ*_*i*_ is the vector of white noises and it follows a bivariate normal distribution with mean *µϵ* = [0, 0] and covariance matrix Σ_ϵ_ (*ϵ*_*i*_ ∼ *N* (0, Σ_ϵ_)).

The mean level *m*_*i*_ does not change for long intervals and its duration follows a geometric distribution: the probability that *m*_*i*_ takes a new value at any point *i* is regulated by the parameter *η* and when it changes, *m*_*i*_ is incremented by the normal random variable 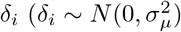, see Supplemental Material for more details).

### PoreMeth2

PoreMeth2 is an R package for the identification of DMRs from Nanopore methylation data of paired samples. It takes as input the methylation calls inferred by tools such as Nanopolish, Guppy or Dorado from a pair of test and matched normal samples and automatically identifies statistically significant DMRs.

DMRs identification is performed by simultaneously segmenting Δ*β* and Δ*S* values of each CpG dinucleotides by using the BiSLM algorithm. DMRs can then be automatically annotated to genic (promoter, introns, exons) and regulatory features (CGIs, Enhancers, TFBS and DHS) to evaluate their functional impact.

The genic elements of PoreMeth2 were generated by parsing the GENCODE project annotation data (release 46 for GRCh38 and release 19 for GRCh37, https://ftp.ebi.ac.uk/pub/databases/gencode/). Consequently, PoreMeth2 contains the annotation of a large number of possible biotypes that include protein coding genes, long non-coding RNAs, pseudogenes, and small RNAs.

For each gene/transcript, PoreMeth2 considers the longest transcript with the highest number of exons and the gene model annotations include Promoter (1kb upstream of the TSS), first exons, internal exons, internal introns and last exons. CpG islands, DHSs, TFBS genomic coordinates were downloaded from UCSC table browser (https://genome.ucsc.edu/cgi-bin/hgTables). Enhancers coordinates were downloaded from https://fantom.gsc.riken.jp/5/.

PoreMeth2 also contains functions to evaluate the quality of methylation data generated by Nanopore sequencing. Given the importance of coverage and data quality to obtain a high resolution in DMRs detection - as previously discussed - we implemented two functions to help visualize statistics about the input data.

The function PoreMeth2SingleExpQualityPlot returns four plots representing the distribution of *β* and *S* values across CpGs genomic positions in a sample and the distribution of the number of reads used to calculate them (Supplementary Figure 17).

The function PoreMeth2PairedExpQualityPlot returns two plots representing the distribution of the number of reads used to calculate Δ*β* and Δ*S* for common CpGs between the two samples (Supplementary Figure 18). PoreMeth2 is publicly available at https://github.com/Lab-CoMBINE/PoreMeth2.

### AML samples sequencing and data preparation

DNA from each of three pairs of matched AMLs was sequenced and basecalled as in [5] and aligned against the human reference genome (hg19) with minimap2. 5mC were inferred with Nanopolish (v. 0.8.5) [21] by using log likelihood ratios (<− 2 or > 2, as suggested in GitHub). RNA from each of three pairs of matched AMLs was sequenced as in [5] with Illumina Novaseq 6000 platform. Transcripts counts from paired-end reads were performed with Salmon v. 0.14.1 and the reference transcriptome GRCh37 from Ensembl. Normalization and statistical analysis were performed with DESeq2 (v. 1.30.1). DEGs with adjusted p-value< 0.05 - scored by Benjamin-Hopkins formula - and absolute *log*_2_*FC* > 0.5 were selected.

### HPGC samples sequencing and data preparation

DNA libraries from T0 and T48 were sequenced with r9.4.1 ONT flowcells on the P2 Solo ONT instrument with a 72 hours acquisition time for each sequencing run.

We used Guppy (v. 6.5.7) with super high accuracy model to obtain basecalls and modified-basecalls. Alignment to the human reference genome (hg19) has been performed by Guppy itself by means of the integrated minimap2 version (2.24). After basecalling, read level 5mCG likelihoods were extracted using Modkit (v. 0.2.2).

cDNA quadruplicates from RNA extraction were sequenced with r9.4.1 flowcells on the P2 Solo ONT instrument with a 72 hours acquisition time. Basecalling and alignment were performed with Guppy (v. 6.5.7). The featureCounts function from the Bioconductor package Rsubread (v 2.12.3) [22] was used to calculate transcript count matrices, while normalization and differential expression analysis were performed with DESeq2. DEGs with adjusted p-value< 0.05 and absolute *log*_2_*FC* > 0.5 were selected.

### Over-representation analysis (ORA)

Pathways for ORA were selected by the network of ‘PATHWAYS IN CANCER’ of KEGG database and the Oncogenic Signaling Pathways in The Cancer Genome Atlas (TCGA) 62. Gene lists of these pathways were downloaded from https://www.kegg.jp/kegg/download/ (KEGG), COSMIC genes from https://cancer.sanger.ac.uk/cosmic/file_download, GEAR genes from http://gear.comp-sysbio.org, TF from http://regnetworkweb.org/. ORA was performed by using fisher-exact test using the list of all UCSC genes as background.

## Data Availability

Nanopore sequencing data for AML samples have been deposited in FASTQ format in the NCBI Sequence Read Archive under accession number PRJNA879930. RNA-Seq data for AML samples in fastq format have been deposited in the NCBI Sequence Read Archive under accession number PRJNA879971.

Nanopore whole-genome sequencing (WGS) and RNA-Seq data for HPGC have been deposited have been deposited in FASTQ format in the NCBI Sequence Read Archive under accession number PRJNA1160066. RNA-Seq counts and methylation frequency and entropy data are available at Gene Expression Omnibus under accession number GSE277456.

## Acknowledgements

We acknowledge financial support under the National Recovery and Resilience Plan (NRRP), Mission 4, Component 2, Investment 1.1, Call for tender No. 104 published on 2.2.2022 by the Italian Ministry of University and Research (MUR), funded by the European Union – NextGenerationEU– Project Title

Computational Methods for Third Generation Cancer Genomics – CUP B53D23007820006 - Grant Assignment Decree No. n. 970 adopted on 30/06/2023 by the Italian Ministry of Ministry of University and Research (MUR).

Funded by the European Union - Next Generation EU. Views and opinions expressed are however those of the author(s) only and do not necessarily reflect those of the European Union or the European Commission. Neither the European Union nor the European Commission can be held responsible for them. PNRR MUR M4 C2 Inv. 1.5 CUP B83C22003920001 - Spoke 3 - Subproject 14 “A Computational Platform for future diagnostic”.

## Competing interests

Alberto Magi and Gianluca Mattei have received travel funding for presenting at symposia organized by Oxford Nanopore Technologies and for a poster presentation, respectively

