## Supplemental Material for "PoreMeth2: decoding the evolution of methylome alterations with Nanopore sequencing"

### AML Samples Library Preparation and Sequencing

DNA was qualitatively and quantitatively analyzed using Nanodrop and Qubit respectively. The sequencing was performed on the GridION 5X platform (Oxford Nanopore Technologies, Oxford, UK). Library preparation was performed according to the Oxford Nanopore Technologies (ONT) manufacturer's protocol for genomic DNA, the Ligation Sequencing Kit 1D SQK-LSK109. For each library 1 $\mu g$ of DNA as starting material was used. The input DNA for all nanopore libraries was unshared. Initially DNA was mixed with 3,5 ul of NEBNext FFPE DNA Repair Buffer and 2 $\mu l$ of NEBNext FFPE DNA Repair Mix (M6630, NEB, Ipswich, Massachusetts) in order to repair possible nicks, and then end-prepped adding 3,5 ul of Ultra II End-prep reaction buffer and 3 $\mu l$ of Ultra II End-prep enzyme mix (E7546, NEB). The reaction was incubated at 20°C for 5 minutes and 65°C for 5 minutes, followed by a purification step with 60 $\mu l$ of 1X AMPure XP beads, by a washing and an elution. The end-repaired DNA was ligated with 5 $\mu l$ Adapter Mix (AMX, ONT) using 10 $\mu l$ NEBNext Quick T4 DNA Ligase (E6056, NEB) for 10 min at room temperature. The adapted-ligated DNA was cleaned up by adding 40 $\mu l$ of 1X AMPure XP beads and incubated at room temperature for 5 min. The beads were pelleted on a magnetic rack and pellet was washed twice by resuspending in 250 $\mu l$ Long Fragment Buffer (LFB, ONT) in order to enrich for long molecules. Then, the reaction was eluted in 15 $\mu l$ of Eluition Buffer (EB, ONT) at room temperature for 10 minutes. The prepared library was quantified by Qubit and then, prior to loading onto the flow cell, was mixed with 37,5 $\mu l$ of Sequencing Buffer (SB, ONT) and 25,5 $\mu l$ of Loading Beads (LB, ONT). According to the ONT's protocol before loading, each R9.4.1 SpotON flow cell (FLO-MIN106D) was primed mixing the Flush Tether (FLT, ONT) directly to the tube of Flush Buffer (FB, ONT) and loading this mix via the priming port. Libraries were sequenced on a GridION 5x device and, in order to maximize the total throughput, sequencing was carried out for 48 h.

### HPGC Samples Library Preparation and Sequencing

Human peripheral glial cells (HPGC) (P10351, Innoprot) were cultured and maintained in dedicated medium (P60123, Innoprot) at 37°C in 5% CO2 and 95% O2. HPGC were cultured in a 6 cm Petri dish until the 80% of confluence and the day after were treated for 0 (T0) or 48 (t48) hours with G protein-coupled receptor agonist (S1198, Merck). Genomic DNA and total RNA were then extracted using DNeasy Blood  Tissue Kit (69506, Qiagen) and RNeasy Mini Kit (74106, Qiagen) respectively. Before Nanopore Library preparation, total RNA was quantified via Qubit Fluorometer with RNA HS

Assay Kit (32852, Thermo Fisher Scientific) and RNA quality was validated with Agilent 2100 Bioanalyzer using RNA 6000 Nano Kit (5067-1511, Agilent). For RNA and DNA library preparation, the PCR-cDNA Barcoding Kit (SQK-PCB111.24) and SQK-LSK109-XL kit respectively were used following manufacturer's instructions.

### Bivariate SLM

For DMRs identification we developed a bivariate version of the SLM algorithm.

$\Delta\beta$ and $\Delta S$ signals for N CpGs are modeled as a sequential processes $x = (x_1, .., x_i, .., x_N)$ (with $x_i$=$(\Delta\beta_i, \Delta S_i)$) as the sum of two independent stochastic processes:

$$x_i = m_i + \epsilon_i \quad (1)$$

$$m_i = (1 - z_i) \cdot m_{i-1} + z_{i-1} \cdot (\mu + \delta_i) \quad (2)$$

where :

- $m_i = (m_{i1}, m_{i2})$ is the vector of the unobserved mean level,
- $\epsilon_i$ is the vector of white noises and it follows a bivariate normal distribution with mean $\mu\epsilon = [0, 0]$ and covariance matrix $\sum_\epsilon$ ( $\epsilon_i \sim N(0, \Sigma_\epsilon)$),
- $z_i$ are random variables taking the values in [0,1] with probabilities ($\eta$ = Pr($z_i$ = 1), $(1 - \eta)$ = $Pr(z_i = 0)$,
- $\delta_i$ are random vectors that follow a bivariate normal distribution with mean $\mu$ and covariance matrix $\Sigma_\mu$.

Moreover, $m_i$ depends by $z_i$: when $z_{i-1} = 0$, $m_i$ is the same as $m_{i-1}$ and when $z_{i-1} = 1$, $m_i$ takes its new value according to a bivariate Gaussian law with mean $\mu$ and covariance matrix $\sum_\epsilon$ independently of $m_{i-1}$ :

As $x_i$ are modeled as the sum of two independent stochastic processes, the expected value of $x_i$ is $\mu$, and its covariance matrix is the sum of the covariances of the two processes:

$E(x_i) = \mu = (\mu_{\Delta\beta}, \mu_{\Delta S})$ (3)

$Cov(x_i) = \Sigma_\mu + \Sigma_\epsilon$ (4)

Hence, we can introduce a different parametrization of the SLM by defining the parameter $\omega$ such that $\Sigma_\mu = \omega \cdot \Sigma$ and $\Sigma_\epsilon = (1–\omega) \cdot \Sigma$.

We also hypothesize that the white noise distributions and mean level distributions can be considered independent between $\Delta\beta$ and $\Delta S$ signals:

$N(0,\Sigma_\epsilon) = N(0, \sigma_{\epsilon_{\Delta\beta}}) \cdot N(0, \sigma_{\epsilon_{\Delta S}})$ (5)

$N(\mu, \Sigma_\mu) = N(\mu_{\Delta\beta}, \sigma_{\mu_{\Delta\beta}}) \cdot N(\mu_{\Delta S}, \sigma_{\mu_{\Delta S}})$ (6)

where $\sigma_{\epsilon_{\Delta\beta}}$ and $\sigma_{\epsilon_{\Delta S}}$ are the standard deviations of the white noises of $\Delta\beta$ and $\Delta S$ signals, while $\sigma_{\mu_{\Delta\beta}}$ and $\sigma_{\mu_{\Delta S}}$ are the standard deviations of the mean level distributions.

With these assumptions, the random process $z_i$ is the only variable correlating the two signals. When $z_i$ changes its value, the mean level of $\Delta\beta$ and $\Delta S$ shifts, allowing the joint model to detect common shifts in the mean [**?** ].

As a consequence of the previous definitions, the joint distribution of the observations and latent variables, given the parameters, has this form:

$$\begin{aligned} p(x,m,z|\theta) &= p(x|m,\Sigma_\epsilon)\cdot p(m|z,\mu,\Sigma_\mu)\cdot p(z|\eta) \\ &= \prod_{i=1}^{N} p(x_i|m_i,\Sigma_\epsilon)\cdot p(m_0)\cdot \prod_{i=0}^{N-1} p(m_{i+1}|m_i,z_i,\mu,\Sigma_\mu)\cdot p(z_i|\eta) \end{aligned} \tag{7}$$

where $\Theta = (\mu, \Sigma_\mu, \Sigma_\epsilon, \eta)$ and:

- $p(x_i|m_i,\Sigma_\epsilon) = \mathcal{N}(x_i|m_i,\Sigma_\epsilon)$ is the probability distribution of $x_i$ given $m_i$ and the parameters.
- $p(m_{i+1}|m_i,z_i,\mu,\Sigma_\mu) = (1-z_i)\cdot\delta(m_{i+1}-m_i) + z_i\cdot\mathcal{N}(m_{i+1}|\mu,\Sigma_\mu)$ is the probability distribution of the latent variable $m_{i+1}$ given $m_i$, $z_i$, and the parameters; $\delta$ is the Dirac delta function.
- $p(z_i|\eta) = \eta\cdot\delta(z_i-1) + (1-\eta)\cdot\delta(z_i)$ is the probability density function of $z_i$.

Equation 7 defines a Hidden Markov Model (HMM) of order one, in which a single state variable, $q_i = (m_i, z_i)$, summarizes all the relevant past information of the underlying process. In the model defined by SM1 the elements of the HMM are the following:

- The state transition probability distribution is: $p(q_{i+1}|q_i,\theta) = p(m_{i+1}|m_i,z_i,\mu,\Sigma_\mu)\cdot p(z_i|\eta)$
- The emission probability distribution is: $p(x_i|q_i,\theta) = p(x_i|m_i,\Sigma_\epsilon)$
- The initial state probability distribution is: $p(q_0|\theta) = p(m_0|\mu,\Sigma_\mu)$

### Bivariate SLM algorithm

The joint distribution of Equation 7 defines an Hidden Markov Model (HMM) of order one with state variable $q_i = (m_i, z_i)$ and bivariate emission probability. The fact that the multivariate SLM is an HMM allows us to make use of the several algorithms developed for these kinds of models.

To maximize the likelihood of the multivariate extension of shifting level model we use a procedure similar to that used in [1]. We introduce a markovian stochastic process $s_1,...,s_k$ taking values in $S = 1,2,...,K$. We assume that the conditional probability of $x_i$, given $s_i = k$, is a bivariate truncated normal with mean $\mu_k = (\mu_{1k}, \mu_{2k})$ and variance $\Sigma_\epsilon$, and the parameter $\mu_k$ is associated to each state of the markovian stochastic process and represents an approximation of the $m_i$ latent variables of the bivariate SLM.

Remembering equations 7 and 6 and that $\Delta\beta$ and $\Delta S$ take values in the range [-1,1], we assume that the conditional probability of $x_i$, given $s_i = k$ is a truncated gaussian distributions with upper and lower bound 1 and -1 respectively, the emission probability distribution has the following form:

$$f_k(x) = \frac{1}{\sqrt{2\pi}\sigma_\epsilon}\exp\left[-\frac{1}{2}\left(\frac{x-\mu_k}{\sigma_\epsilon}\right)^2\right]\cdot\frac{1}{\Phi(\frac{1-\mu_k}{\sigma_\epsilon})-\Phi(\frac{-1-\mu_k}{\sigma_\epsilon})}. \tag{8}$$

where $\Phi$ represent the cumulative distribution functions of a non-truncated Gaussian of parameter $\mu_k$ and $\sigma_\epsilon$.

To complete the description of the model, it remains to specify the state transition matrix $P$. From (**??**) the state transition probability has the following form:

$$\begin{aligned} p(m_{i+1}|m_i,z_i,\mu,\sigma^2,\omega)\cdot p(z_i|\eta) \quad = \quad & [(1-z_i)\cdot\delta(m_{i+1}-m_i) + z_i\cdot N(m_{i+1}|\mu,\omega\cdot\sigma^2)]\cdot \\ & \cdot[\eta\cdot\delta(z_i-1) + (1-\eta)\cdot\delta(z_i)], \end{aligned} \tag{9}$$

hence the state transition matrix is:

$$P_{jk} = \begin{cases} (1-\eta)+\eta\cdot g_{jk} & j = k \\ \eta\cdot g_{jk} & j \neq k \end{cases} \tag{10}$$

where

$$
\begin{aligned}
g_{jk} &= c_j \cdot e^{-\frac{(\mu_k-\mu)^2}{2\sigma_\mu^2}}, \\
c_j &= \left( \sum_{k=1}^{K} e^{-\frac{(\mu_k-\mu)^2}{2\sigma_\mu^2}} \right)^{-1}.
\end{aligned}
\tag{11}
$$

To estimate the parameters of the truncated-gaussian bivariate Shifting Level Model (BiSLM), we develop a two-step algorithm that follows our previous idea in Magi *et al.* [2]. Since the$\Delta\beta$ and $\Delta S$ can take values in a well-defined range, we used a large number of states $K_0$ and we choose $\mu_k = (\mu_{k2}, \mu_{k2})$ in order to densely and homogeneously cover al the combination of the ranges ([-1,1],[-1,1]) instead of estimating the $\mu_k$ parameters by using the Baum and Welch algorithm. Moreover, since we expect that the great majority of CpG have neither differential methylation nor entropy between samples ($\Delta\beta \sim 0$,$\Delta S \sim 0$), we initialize $\mu = (0, 0)$. This simple solution drastically improves the computational performance of our algorithm without affecting its accuracy in the detection of signal shifts.
In the first step of the algorithm we initialize the mean $\mu$ and the variances $\sigma^2$, $\sigma_\mu^2$ and $\sigma_\epsilon^2$ with the following formulas:

$$
\begin{aligned}
&\mu = (0, 0), \\
&\sigma = \sqrt{\tfrac{\sum_{i=1}^{N}(x_i-\mu)^2}{(N-1)}}, \\
&\sigma_\mu^2 = \omega \cdot \sigma^2, \\
&\sigma_\epsilon^2 = (1-\omega) \cdot \sigma^2.
\end{aligned}
\tag{12}
$$

In the second step we apply the Viterbi algorithm to find the best state sequence $s^{(j)}$ and estimate the points of mean shift $z_i$. After Viterbi algorithm we calculate the median of the $\Delta\beta$ and $\Delta S$ values that belong to each segment.

The inputs to the algorithm are the $\Delta\beta$ and $\Delta S$ values ($\Delta\beta = (\Delta\beta_1, .., \Delta\beta_i, .., \Delta\beta_N)$, $\Delta S = (\Delta S_1, .., \Delta S_i, .., \Delta S_N)$) to be segmented, number of states $K_0$, the parameter $\omega$ and the parameter $\theta$.

### Synthetic Data Generation

Synthetic methylation profiles were simulated by first generating long methylation patterns, made of N CpG dinucleotides, as a sequence of consecutive short epialleles (made of three, see Supplemental Figure 1) randomly sampled from multinomial probability distributions that mimic methylated and unmethylated epialleles. The multinomial distribution models the probability of counts of the eight possible three-CpG epialleles ('111','110','101','') with the parameter $p = (p_1, p_2, p_3, p_4, p_5, p_6, p_7, p_8)$. By using multinomial probability distributions with different values of $p$ it is possible to sample methylated or unmethylated epialleles with different level of randomness in different position of the pattern (see Figure S1).

To mimic real methylation patterns, mainly made of methylated CpGs, consecutive short epialleles were sampled from multinomial distribution with $p = (p_1 = 0.9, p_2 = (1-p_1)/3, p_3 = (1-p_1)/3, p_4 = (1-p_1)/3, p_5 = 0, p_6 = 0, p_7 = 0, p_8 = 0)$. We then added, in specific position of the pattern, consecutive epialleles sampled from multinomial distribution with different values of $p$.

Low-entropy methylated epialleles were generated using $p_5 = p_6 = p_7 = p_8 = 0$, $p_1 > 0.5$ and $p_2 = p_3 = p_4 = (1-p_1)/3$. Similarly, low-entropy unmethylated epialleles were generated using $p_1 = p_2 = p_3 = p_4 = 0$, $p_5 > 0.5$ and $p_6 = p_7 = p_8 = (1-p_5)/3$. High-entropy methylated epialleles are generated with $p_5 = p_6 = p_7 = p_8 = 0$ and $p_1 = p_2 = p_3 = p_4 = 0.25$, while high-entropy unmethylated epialleles are generated with $p_5 = p_6 = p_7 = p_8 = 0.25$ and $p_1 = p_2 = p_3 = p_4 = 0$ (see Figure S1.a).

By following these strategy we generated M methylation patterns for test and control samples that allow to simulate methylated or unmethylated epialleles with a predefined level of randomness (entropy). In particular we generated consecutive epialleles made of 5-50 CpGs (5, 10, 15, 20, 25, 30, 35, 40, 45 and 50 CpGs) that simulate hyper- or hypo-methylation (between test and control) with hyper-, hypo- or iso-entropic changes (see Figure S1.b).

The M methylation patterns were finally used to generate long reads by sampling segments of consecutive CpGs from the simulated patterns in order to obtain the desired sequencing coverage (see Figure

S1.c). To test the performance of BiSLM we simulated methylation patterns with epiallelic changes of different sizes (from five to 50 CpGs) and we sampled segments to generate sequencing coverages from 10x to 50x.

#### Synthetic Data Analysis with BiSLM

In order to evaluate the ability of our BiSLM to identify DMRs of different size and with different epiallelic changes we simulated synthetic methylation profiles. To this end, we developed a computational recipe that simulates long methylation patterns, made of N CpG dinucleotides, by generating consecutive short epialleles randomly sampled from a multinomial probability distribution (see Methods and Supplemental Figure S2). Methylation patterns are then exploited to generate long reads datasets with desired sequence size and sequencing coverage.

By using this strategy we simulated test and control samples at sequencing coverages that range from 10x to 50x (10, 20, 30, 40 and 50x) with hypo- and hyper-methylated DMRs from 5 to 50 CpGs (5, 10, 15, 20, 25, 30, 35, 40, 45, 50). hypo- and hyper-methylation was simulated with random and non-random epiallelic shifts and different level of randomness (different multinomial probability distribution, see caption of Figure S2).

For each pair of test and control samples we calculated frequency $\beta$ and entropy $S$ at each CpG and we then applied BiSLM to simultaneously segment $\Delta\beta$ and $\Delta S$. A simulated DMR is considered a true positive if it has a reciprocal overlap larger than 0.9 with a region detected by BiSLM.

As a first step we exploited synthetic data to evaluate the performance of our approach as a function of parameter settings ($\eta$ and $\omega$). Supplemental Figures S3-7 show that the best F-measure (trade off between precision and recall) is obtained for $\omega = 0.1 - 0.2$ and $\eta = 10^{-5} - 10^{-6}$ for all sequencing coverages (10x-50x). As a further step, synthetic data were also used to study the capability of BiSLM to identify DMRs of different size and recognize their epiallelic changes.

The results of our synthetic analysis show (Supplemental Figure S8) that sequencing coverage has a strong effect on the global performance of our segmentation strategy. The identification of DMRs as small as five consecutive CpGs requires at least a sequencing coverage larger than 20x. Large sequencing coverage (30x) is also necessary to correctly classify epiallelic formation. Remarkably, iso-entropic DMRs are more difficult to be identified than hyper- and hypo-entropic alterations.

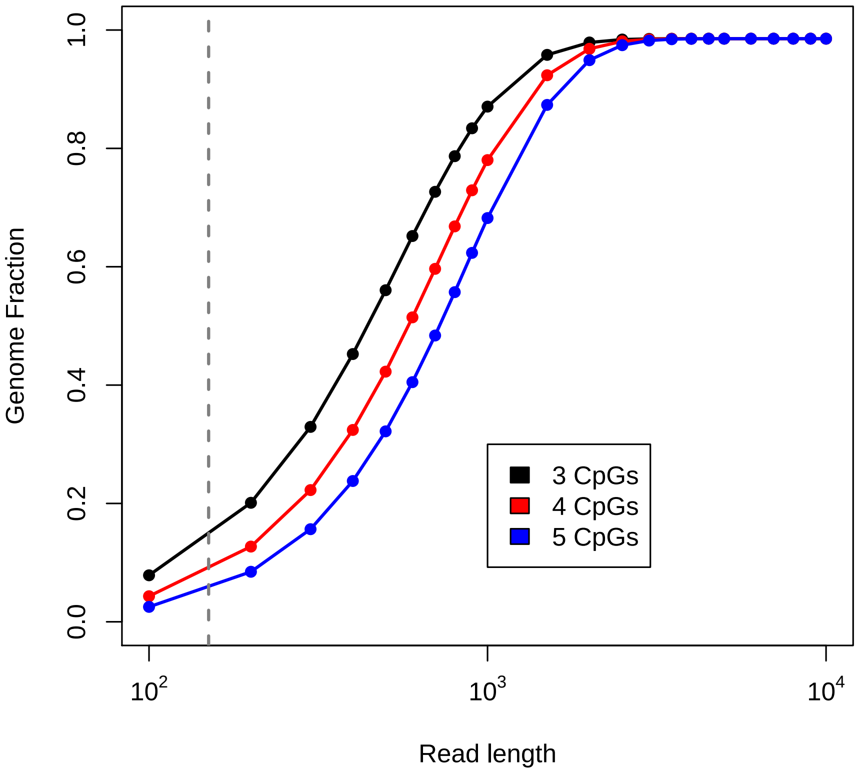


Figure S1: The plot shows the mean coverage across CpG dinucleotides reached by simulated reads of different sizes, spanning from 100 bp to 10 kb. The dotted line represents the mean coverage reached by Illumina sequencing technology, highlighting how Second Generation Sequencing data only allow epialleles characterization in a small portion of the genome (10-15%). The blue, red and black lines display the distribution of the covered genome fraction when applying the bivariate SLM to simulated epialleles with different number of CpGs.

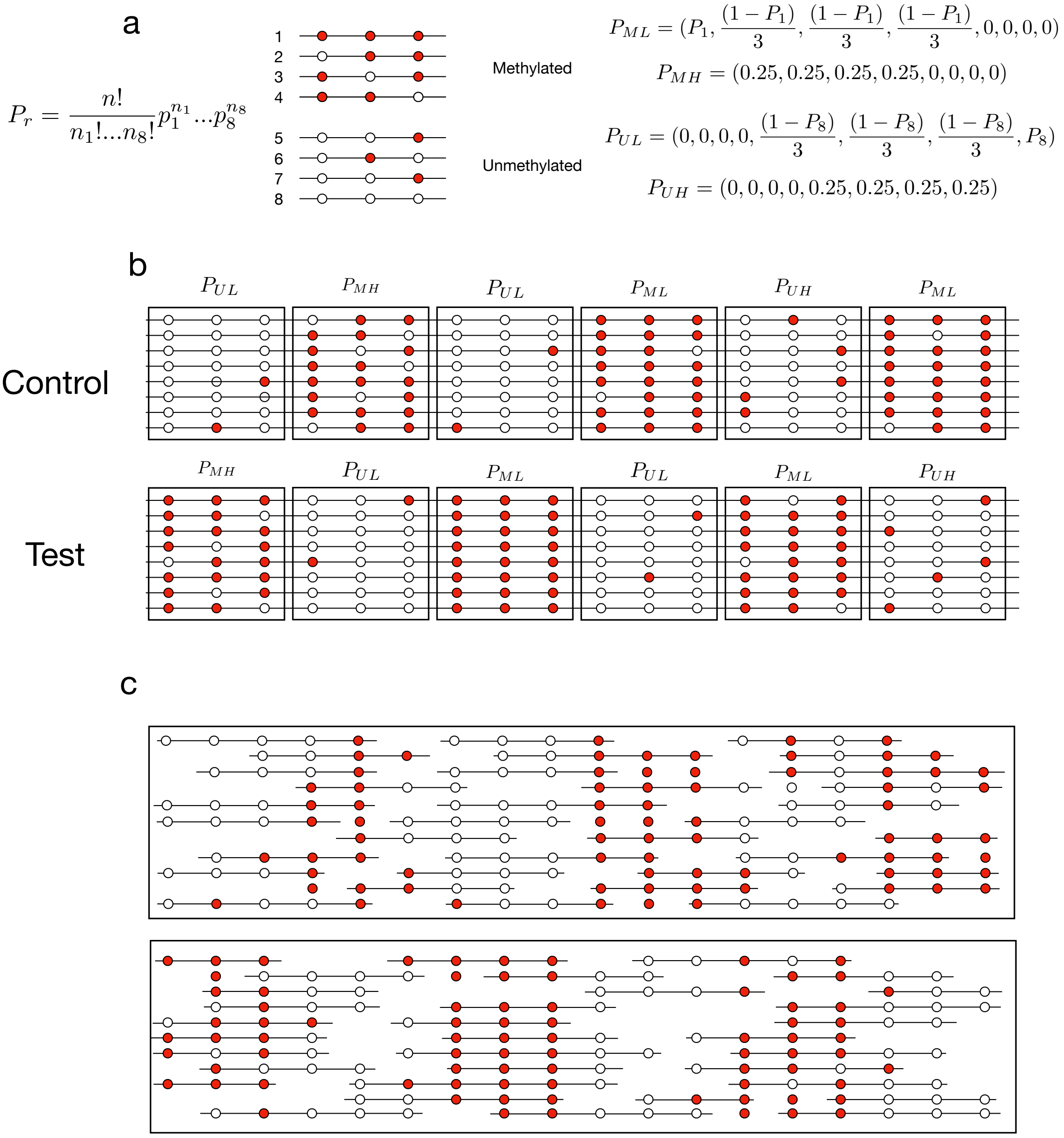


Figure S2: Computational scheme to generate synthetic methylation profiles. Methylation patterns are generated as a sequence of consecutive three-CpGs epialleles ('111','011','101','110') randomly sampled from multinomial probability distribution. To simulate methylated and unmethylated patterns with different level of randomness we used four distributions with parameter $P_{ML}$ (methylated with low stochasticity), $P_{MH}$ (methylated with high stochasticity), $P_{UL}$ (unmethylated with low stochasticity) and $P_{UH}$ (unmethylated with high stochasticity). Setting different values of $P_1$ and $P_8$ allows to modulate the stochasticity of the epialleles and consequently methylation entropy. In our simulations we set $P_1 > 0.5$ and $P_8$ with values ranging between 0.5 and 0.9. Following these scheme we generated methylation patterns for test and control samples that simulate all the six possible epiallelic changes (b). The final step consists in sampling segments of consecutive CpGs from the simulated patterns in order to obtain the desired sequencing coverage. To test the performance of BiSLM we simulated methylation patterns with epiallelic changes of different sizes (from five to 50 CpGs) and we sampled segments to generate sequencing coverages from 10x to 50x.

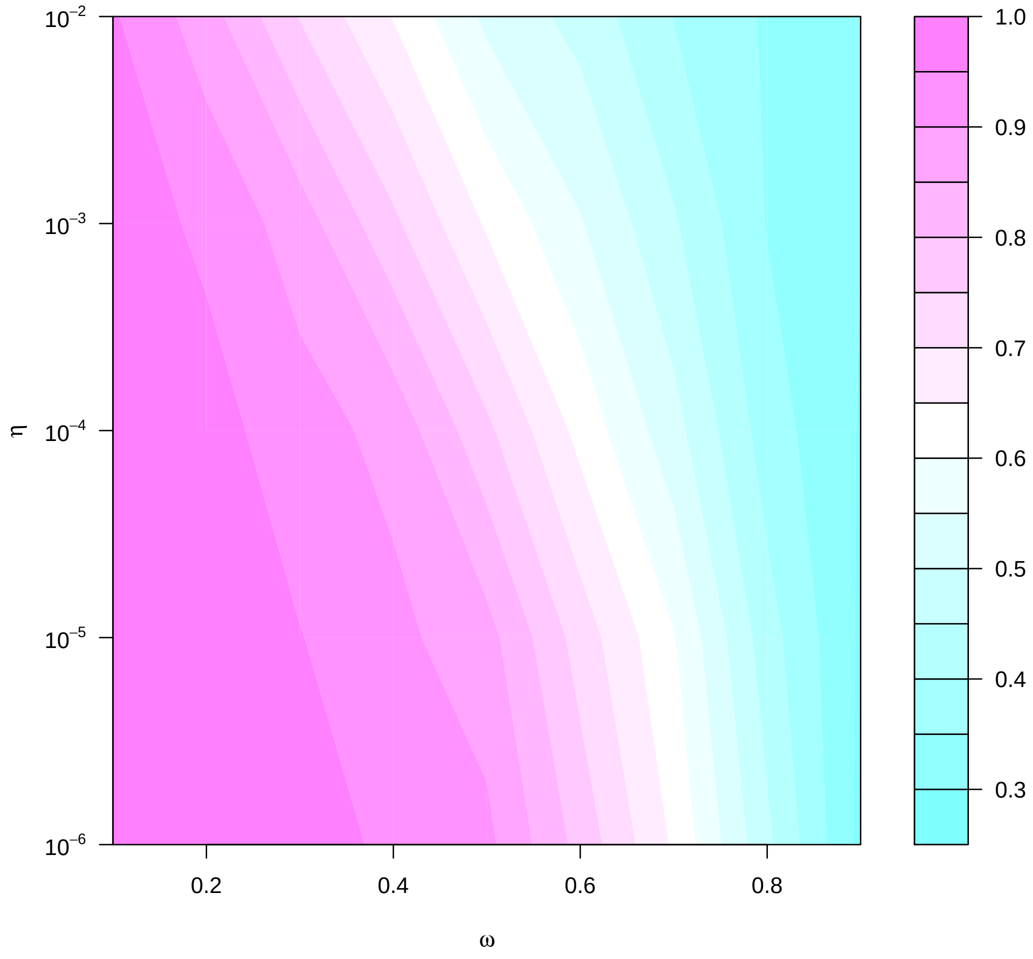


Figure S3: The contour plot shows the performance (in terms of F-Measure) of BiSLM for different combinations of values of $\eta$ and $\omega$ parameters in the analysis of methylation profiles generated from synthetic epialleles at 10x of sequencing coverage.

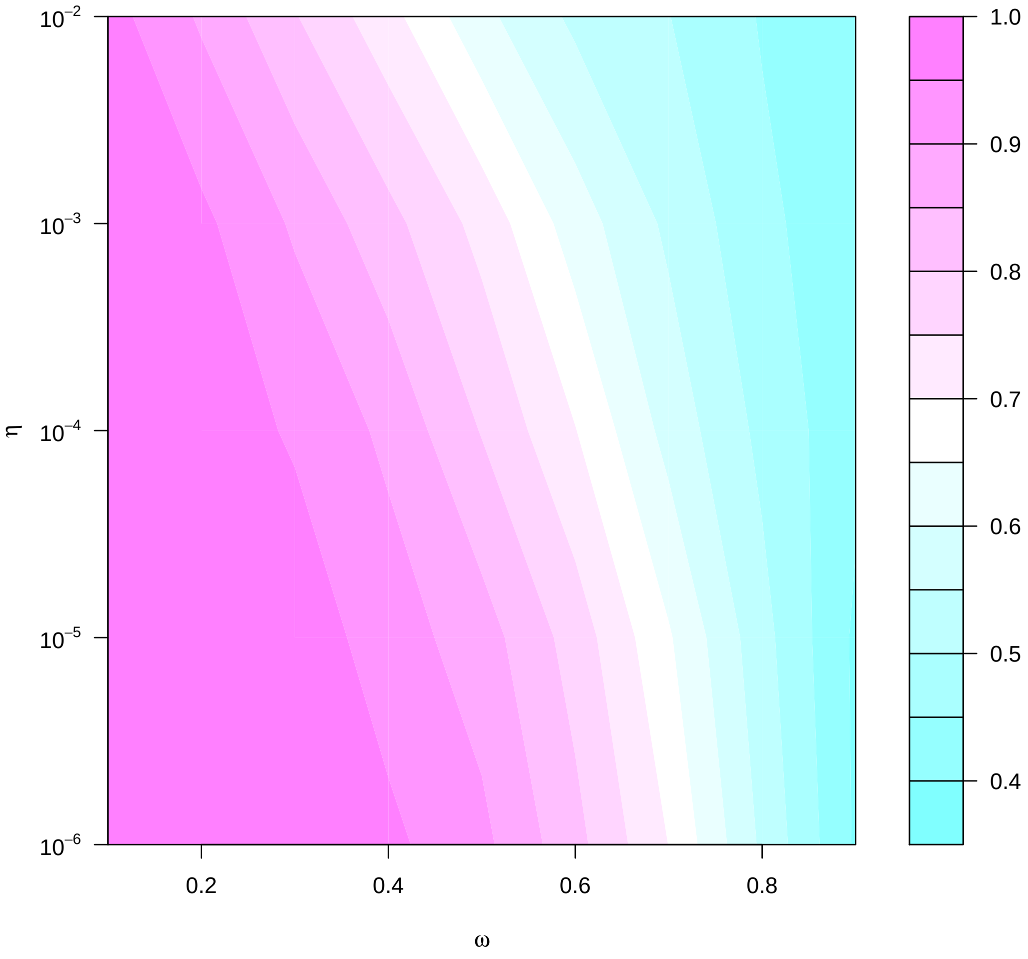


Figure S4: The contour plot shows the performance (in terms of F-Measure) of BiSLM for different combinations of values of $\eta$ and $\omega$ parameters in the analysis of methylation profiles generated from synthetic epialleles at 20x of sequencing coverage.

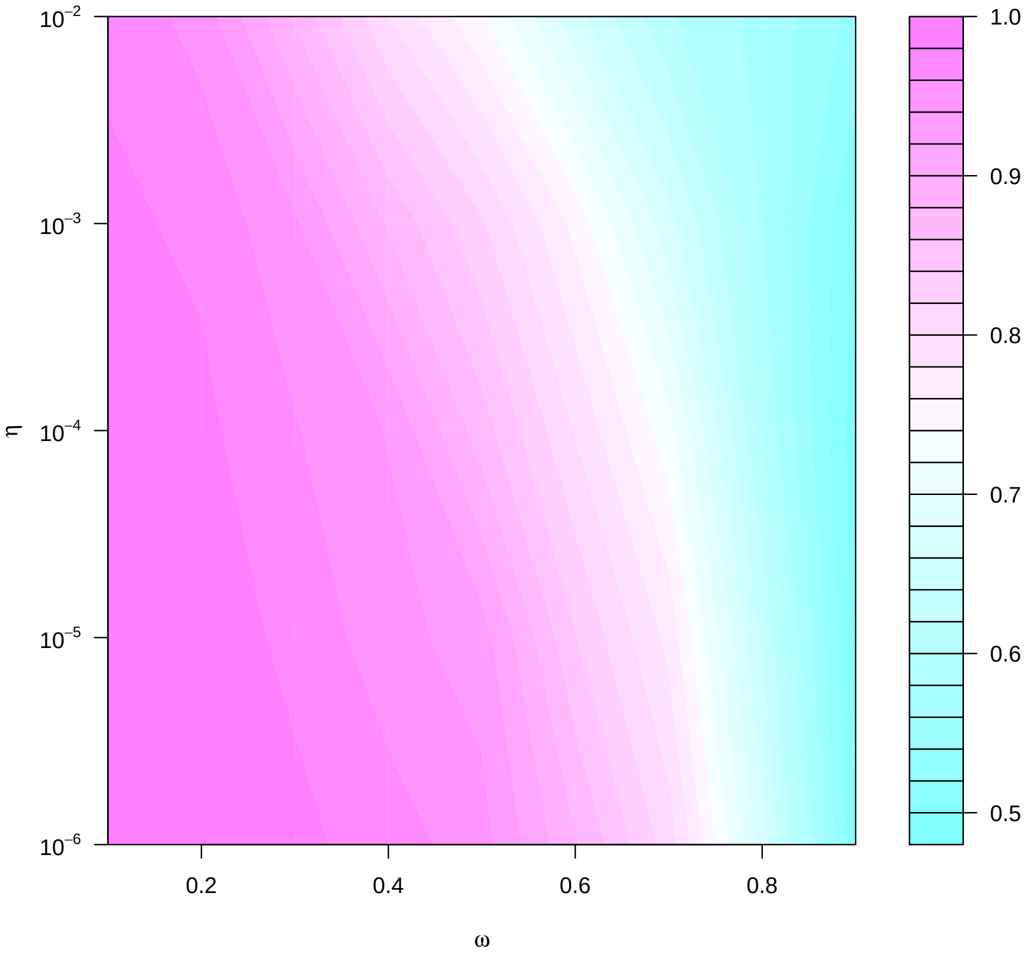


Figure S5: The contour plot shows the performance (in terms of F-Measure) of BiSLM for different combinations of values of $\eta$ and $\omega$ parameters in the analysis of methylation profiles generated from synthetic epialleles at 30x of sequencing coverage.

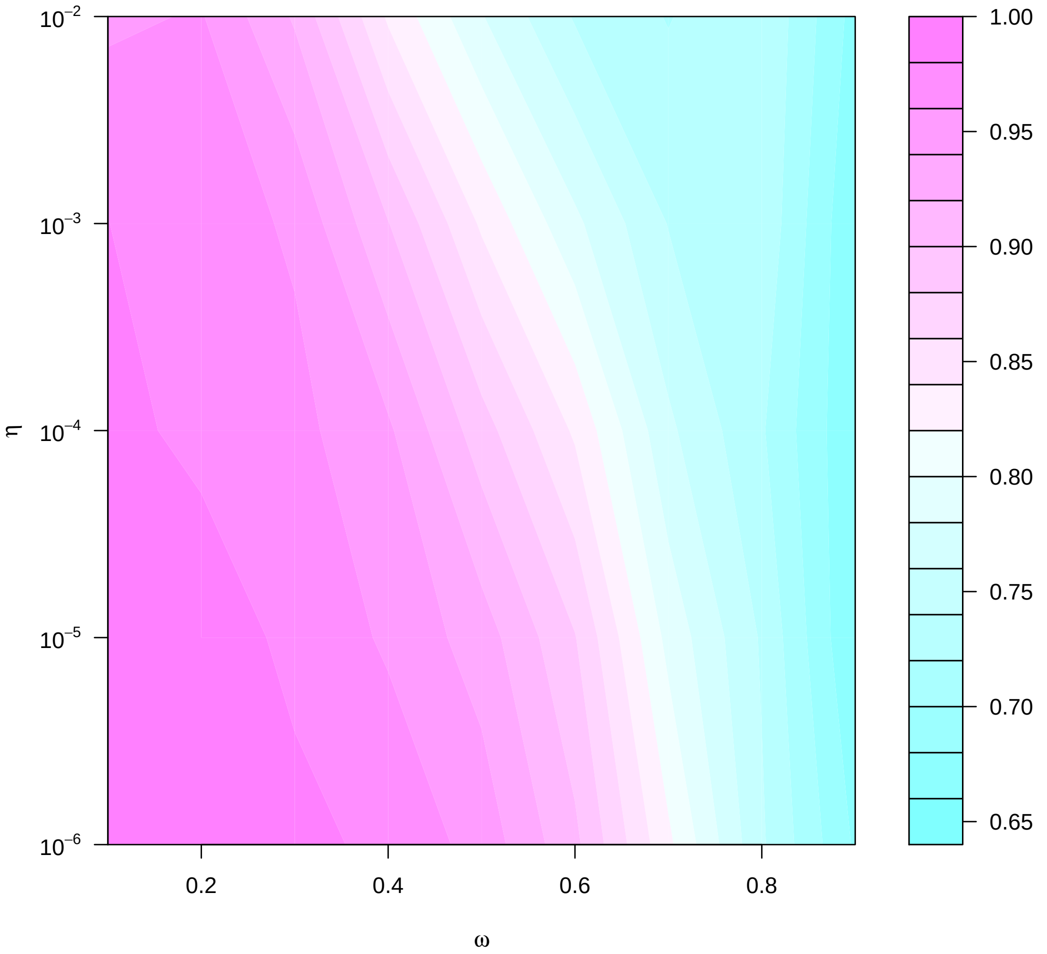


Figure S6: The contour plot shows the performance (in terms of F-Measure) of BiSLM for different combinations of values of $\eta$ and $\omega$ parameters in the analysis of methylation profiles generated from synthetic epialleles at 40x of sequencing coverage.

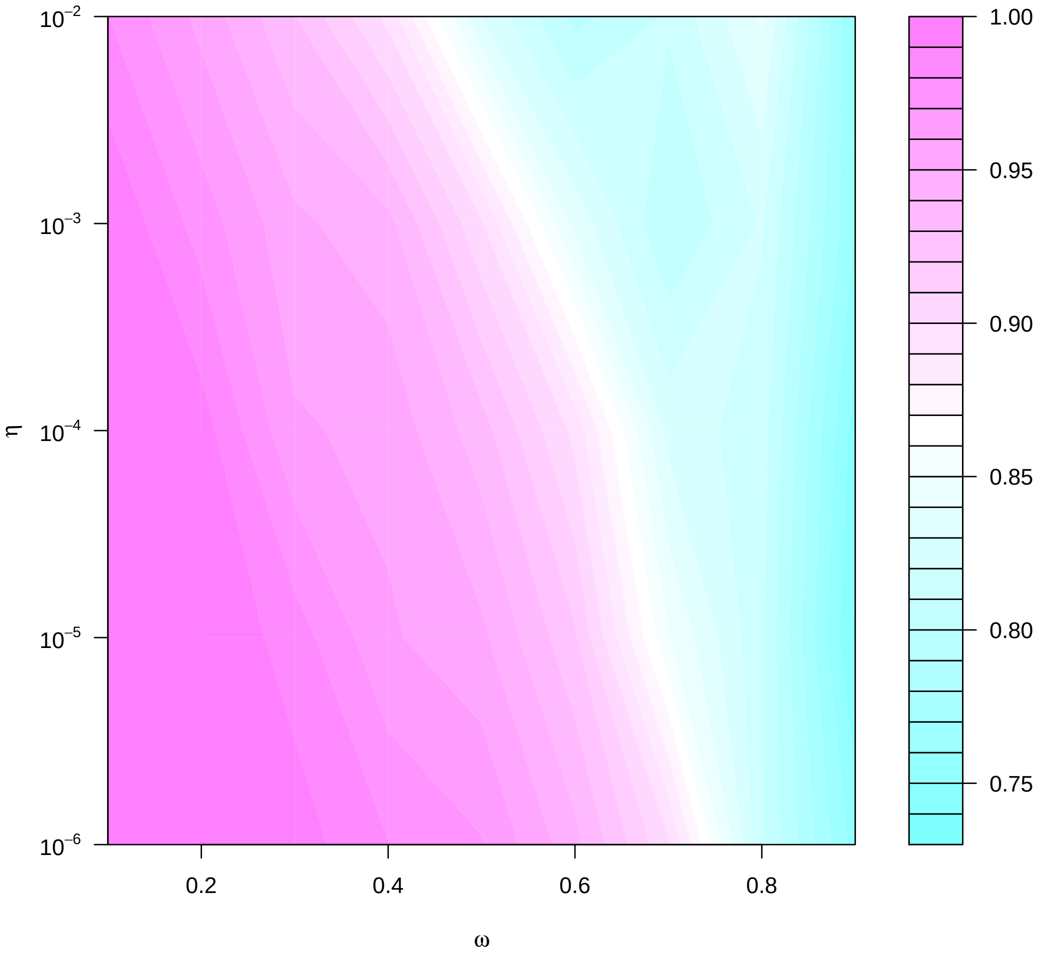


Figure S7: The contour plot shows the performance (in terms of F-Measure) of BiSLM for different combinations of values of $\eta$ and $\omega$ parameters in the analysis of methylation profiles generated from synthetic epialleles at 50x of sequencing coverage.

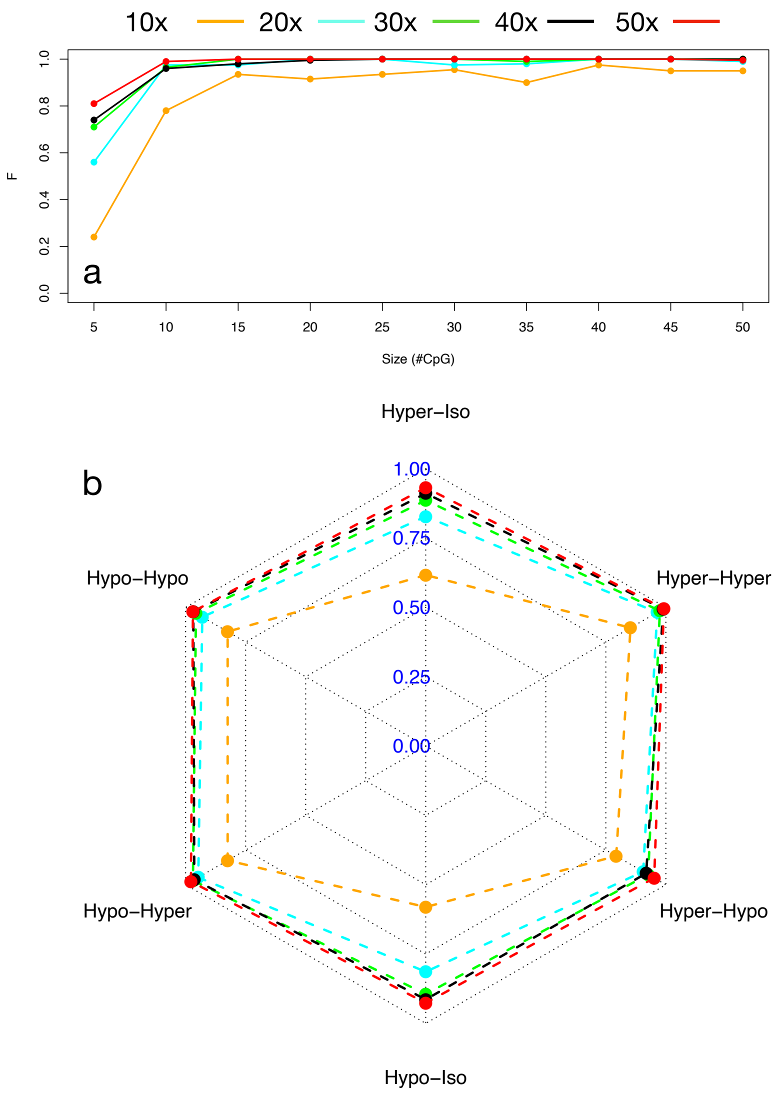


Figure S8: Performance of bivariate SLM on the analysis of synthetic methylation profiles. Panel (a) reports the F-measure (trade-off between precision and recall) obtained by SLM in the detection of synthetic DMRs of different size as a function of sequencing coverage. The radarplot of panel (b) shows the correct classification rate (CCR) obtained by bivariate SLM in correctly classifying different classes of epiallelic shifts for different sequencing coverages.

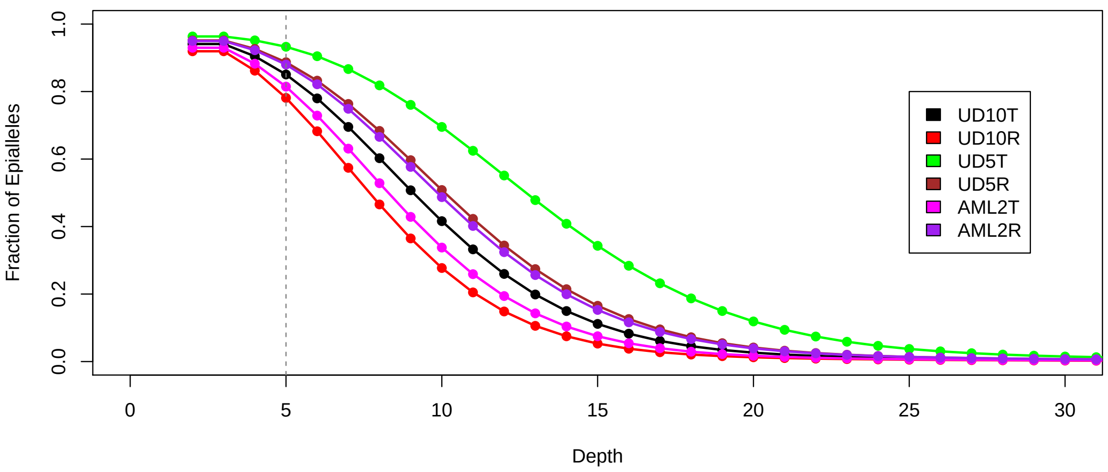


Figure S9: The figure shows the fraction of epialleles as a function of sequencing coverage (depth) for the six AML samples. Vertical dotted line indicates 5x coverage.

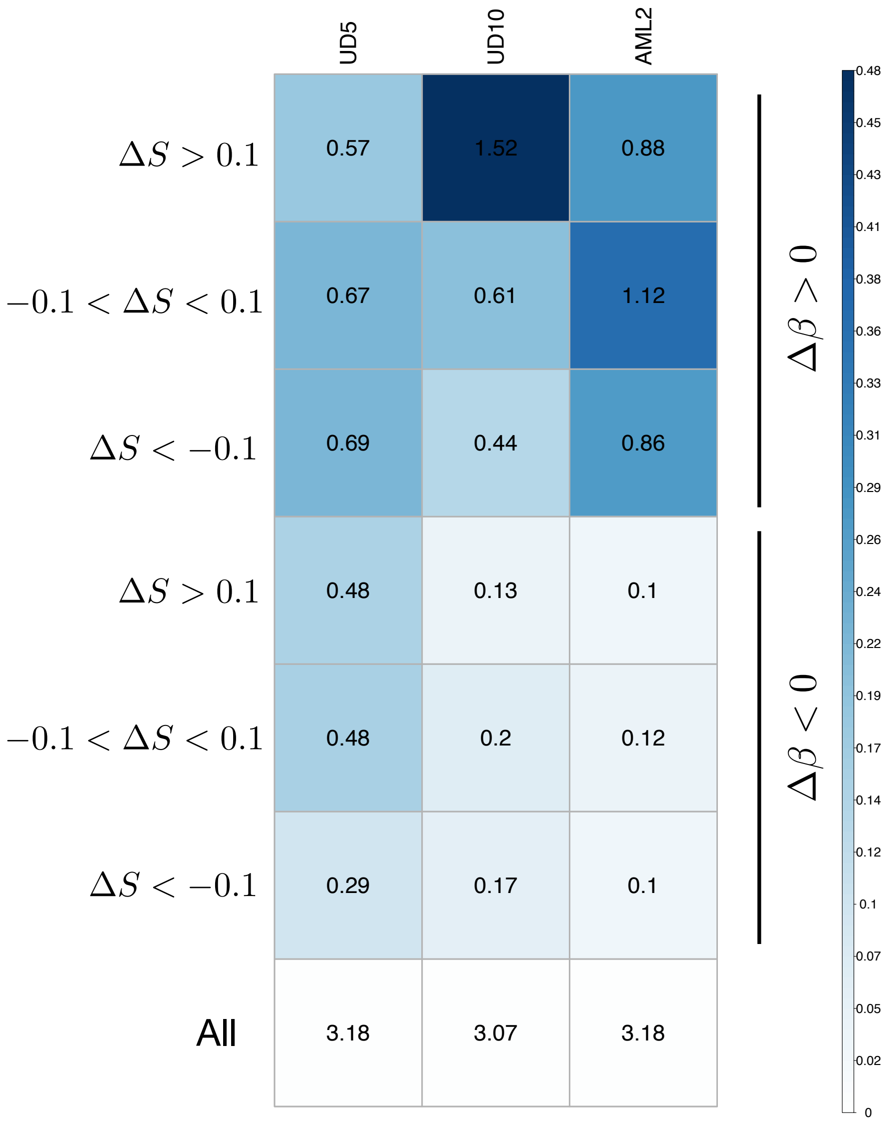


Figure S10: The figure shows the cumulative size of DMRs (in Mb), overall and for different $\Delta S$ and $\Delta\beta$ categories for the three AML sample pairs. While single DMRs show a similar size distribution across the six categories, the cumulative size is significantly higher for hyper-methylated regions ($\Delta\beta > 0.2$) compared to hypo-methylated regions ( $\Delta\beta <$ -0.2)

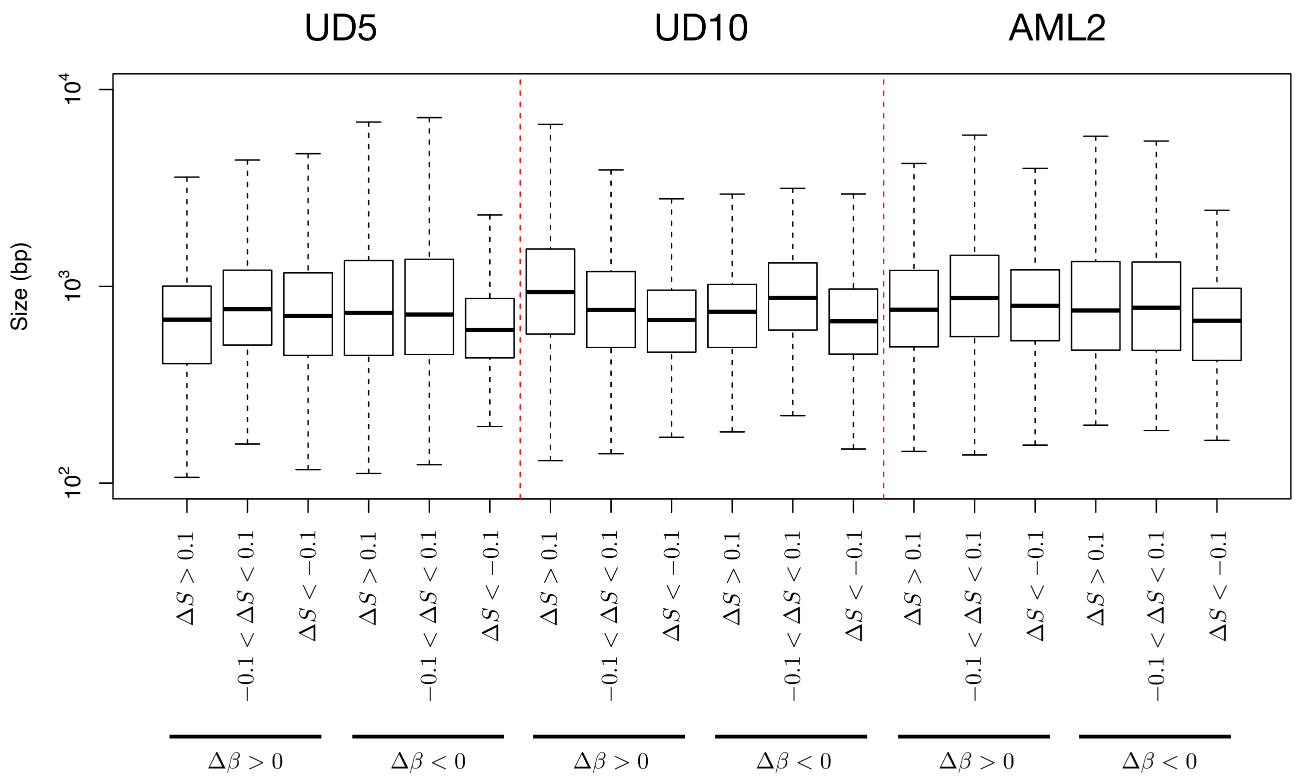


Figure S11: The figure shows DMRs size distribution for different $\Delta S$ and $\Delta\beta$ categories across the three AML sample pairs.

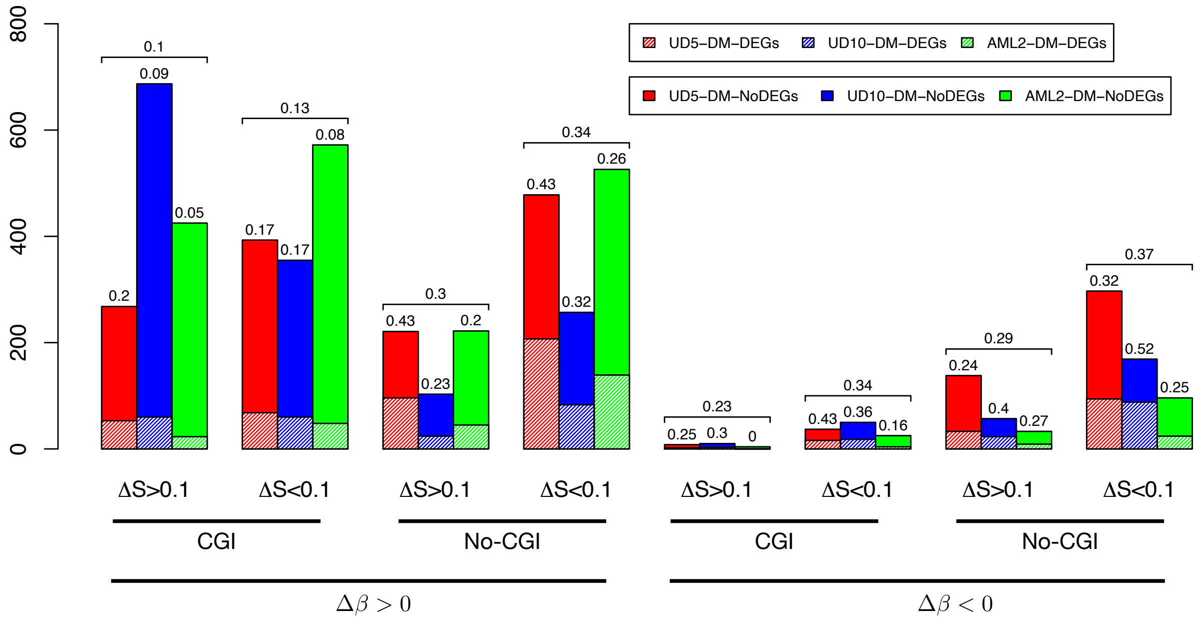


Figure S12: Figure shows the proportion of DMGs that are also DEGs for different $\Delta S$ and $\Delta\beta$ categories for the three AML sample pairs. Data are reported for DMRs overlapping CpG islands (CGI) and outside CpG islands (NoCGI). Textured bars show the number of DM-DEGs.

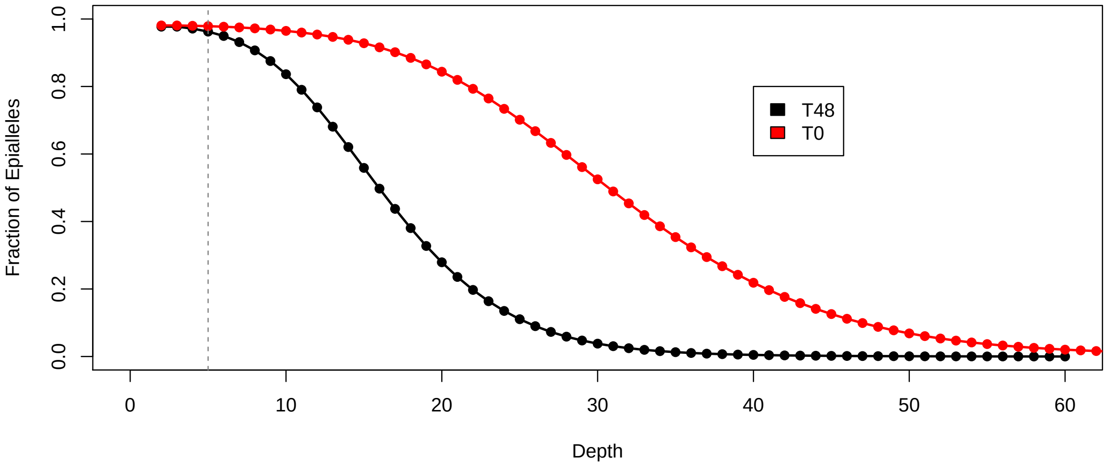


Figure S13: The figure shows the fraction of epialleles as a function of sequencing coverage (depth) for the Shwann dataset. Vertical dotted line indicates 5x coverage.

| | $\Delta\beta > 0$ | | | $\Delta\beta < 0$ | | | |
|---|---|---|---|---|---|---|---|
| | $\Delta S > 0.1$ | $-0.1 < \Delta S < 0.1$ | $\Delta S < -0.1$ | $\Delta S > 0.1$ | $-0.1 < \Delta S < 0.1$ | $\Delta S < -0.1$ | All |
| T48 | 4.16 | 0.55 | 2.73 | 0 | 0.01 | 0.01 | 7.45 |

Figure S14: The figure shows the cumulative size of DMRs (in Megabases), overall and for different $\Delta S$ and $\Delta\beta$ categories for the Shwann dataset. While single DMRs show a similar size distribution across the six categories, the cumulative size is significantly higher for hyper-methylated regions ($\Delta\beta > 0.2$) compared to hypo-methylated regions ( $\Delta\beta <$ -0.2)

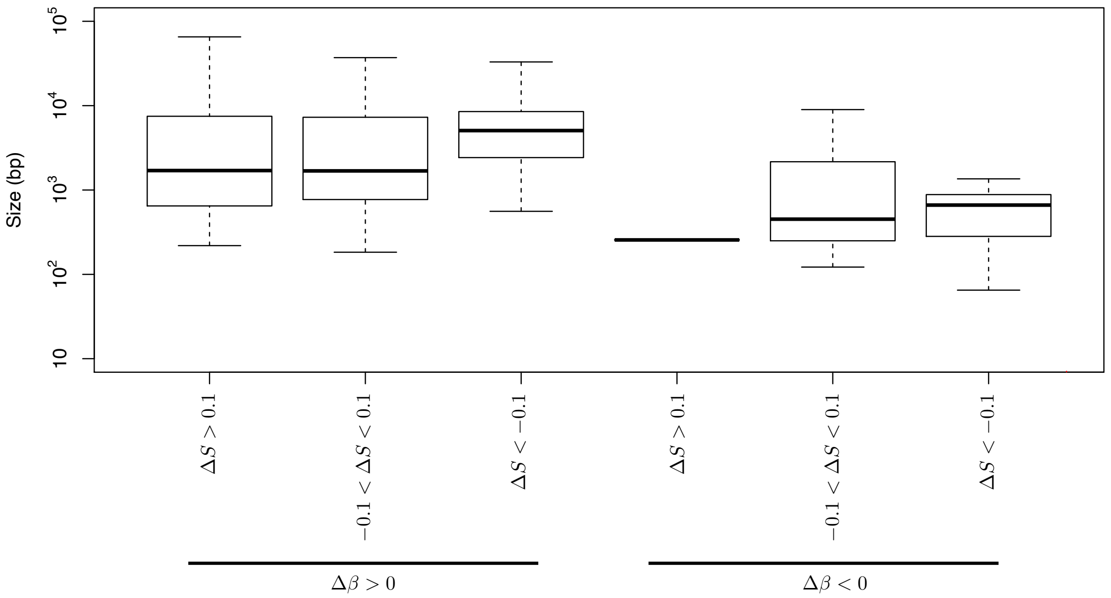


Figure S15: The figure shows DMRs size distribution for different $\Delta S$ and $\Delta\beta$ categories for the Shwann dataset.

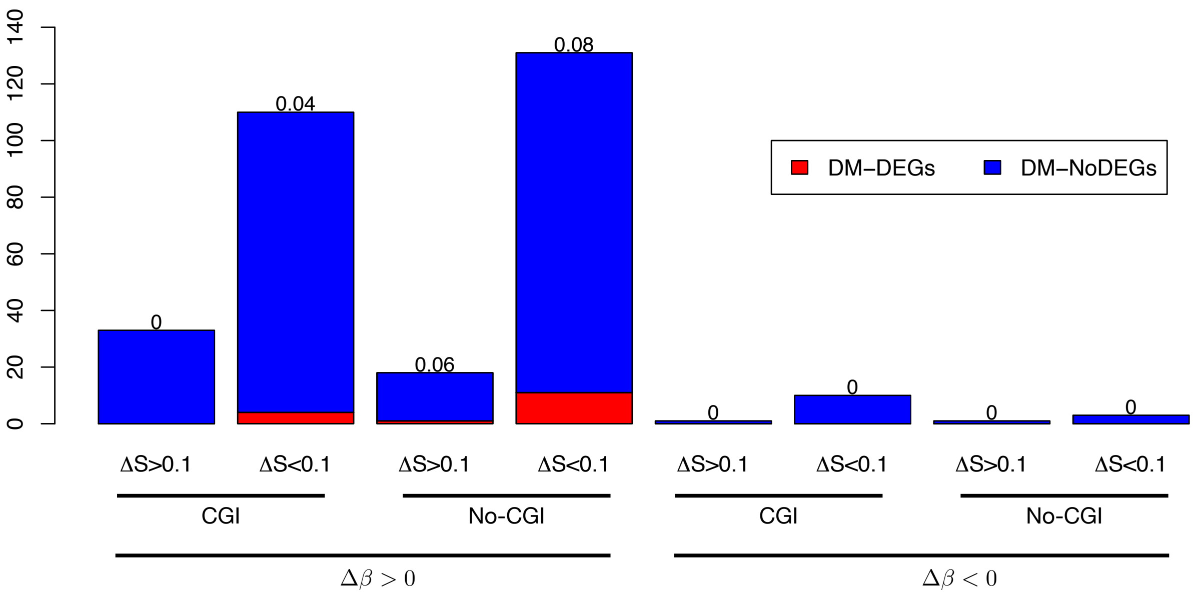


Figure S16: Figure shows the proportion of DMGs that are also DEGs for different $\Delta S$ and $\Delta\beta$ categories for the Shwann dataset. Data are reported for DMRs overlapping CpG islands (CGI) and outside CpG islands (NoCGI). Textured bars show the number of DM-DEGs.

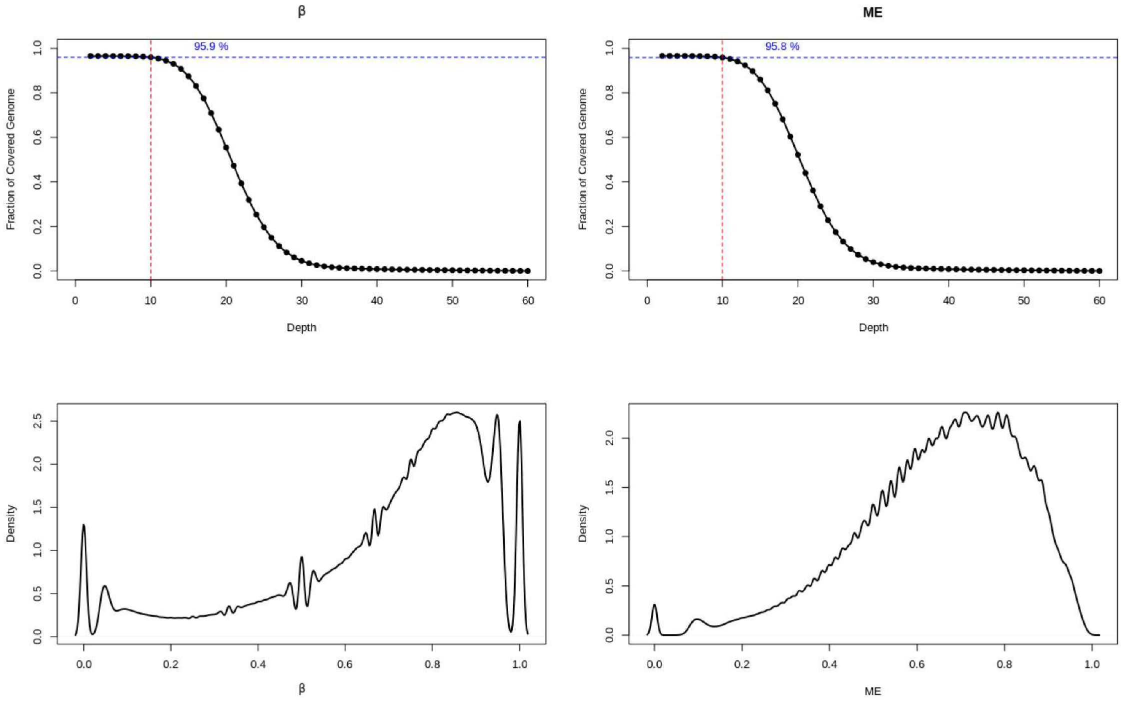


Figure S17: Figure shows an example of Quality Plot generated with PoreMeth2 function PoreMeth2SingleExpQualityPlot. The four quadrants respectively show (from top to bottom and from left to right): i) the distribution of the number of reads (depth) used to calculate $\beta$ in each CpG, ii) the distribution of the number of reads (depth) used to calculate $S$ in each CpG, iii) the distribution of $\beta$ values across the sample and iv) the distribution of $S$ values across the sample. The vertical and horizontal lines in the first two quadrants highlight the fraction of CpGs covered by at least 10 reads.

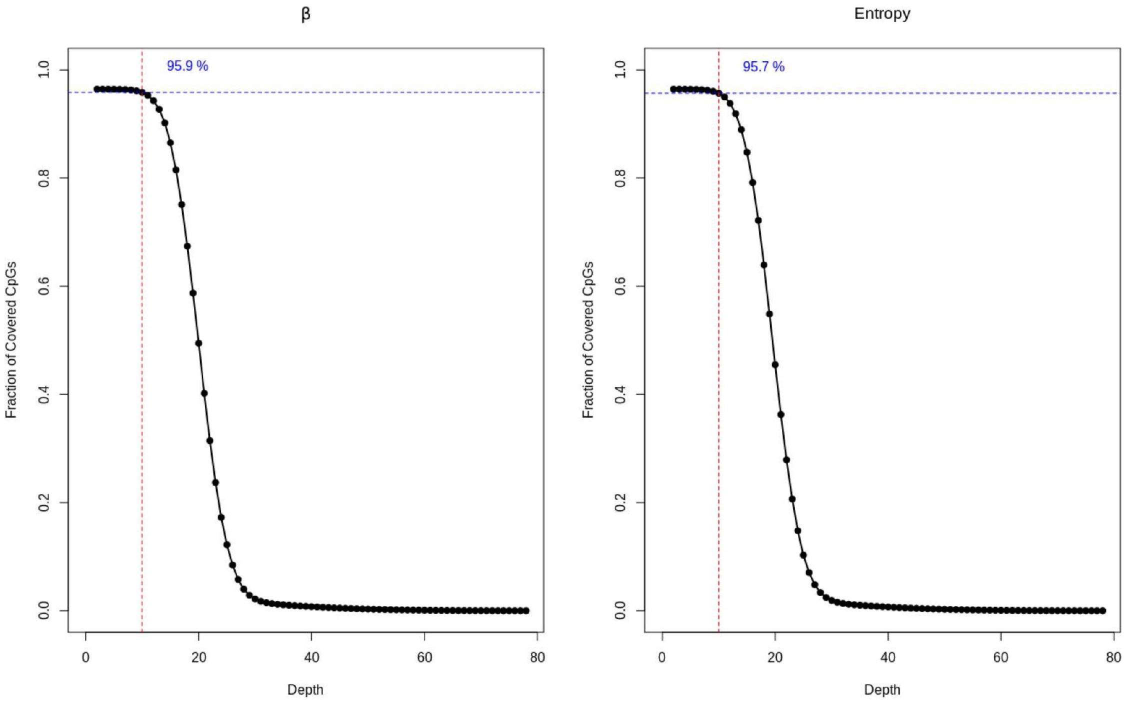


Figure S18: Figure shows an example of Quality Plot generated with PoreMeth2 function PoreMeth2PairedExpQualityPlot. The two quadrants show the distribution of the number of reads (depth) used to calculate $\beta$ and $S$ in each common CpG between the two experiments, with depth being the minimum value among the two samples for each CpG. The vertical and horizontal lines highlight the fraction of CpGs covered by at least 10 reads in both samples.
